# Spatial positioning of preimplantation mouse embryo blastomeres is regulated by mTORC1 and 7mG-cap dependent translation at the 8- to 16-cell transition

**DOI:** 10.1101/2023.03.07.531473

**Authors:** Lenka Gahurova, Jana Tomankova, Pavlina Cerna, Pablo Bora, Michaela Kubickova, Giorgio Virnicchi, Kristina Kovacicova, David Potesil, Pavel Hruska, Zbynek Zdrahal, Martin Anger, Andrej Susor, Alexander W. Bruce

## Abstract

Preimplantation stages of mouse embryo development involve temporal and spatial specification and segregation of three late blastocyst cell lineages; trophectoderm (TE), primitive endoderm (PrE) and epiblast (EPI). Spatial separation of the outer TE lineage from the two inner cell mass (ICM) lineages (PrE and EPI) starts with the 8- to 16-cell transition and concludes following transit through the 16- to 32-cell stages. This results in an early blastocyst ICM derived from descendants of primary founding inner cells and a secondarily contributed population, of which subsequent relative EPI versus PrE potencies are subject to debate. Here, we report generation of primary but not the secondary ICM populations is highly dependent on temporally discreet activation of the mammalian target of Rapamycin (mTOR – specifically mTORC1) during M-phase entry at the 8-cell stage. This role is mediated via regulation of the 7-methylguanosine-(7mG) cap binding initiation complex (EIF4F), linked to translation of a subset of key mRNAs containing 5’ UTR terminal oligopyrimidine (TOP-) or TOP-like sequence motifs; as knockdown of identified TOP-like motif containing transcripts also impairs generation of 16-cell stage primary ICM founders. However, mTOR inhibition induced ICM cell number deficits at the early blastocyst stage can be compensated by the late blastocyst stage, in the absence of inhibition. This compensation is likely initiated at the 32-cell stage when supernumerary outer cells in mTOR-inhibited embryos exhibit molecular characteristics of inner cells. Collectively, the data identify a novel mechanism specifically governing initial spatial segregation of blastomeres in the mouse embryo, that is distinct from those directing subsequent inner cell formation and contributes to germane segregation of late blastocyst lineages.

## INTRODUCTION

Preimplantation stages of mouse embryo development culminate at E4.5 with formation of peri-implantation blastocysts, comprising three distinct cell lineages. These are two differentiating and epithelised lineages, the outer trophectoderm (TE - ultimately contributing to placenta) and the primitive endoderm (PrE – a monolayer residing at the cavity to ICM interface, later forming yolk sac membranes). The third lineage is the pluripotent epiblast (EPI - deep within the ICM, serving as a foetal cell progenitor pool) [1, 2]. Cleavage of apical-basolaterally polarised 8-cell stage blastomeres [3–7] heralds the first relative spatial segregation of embryonic cells. Resultant daughter 16-cell stage blastomeres either occupy outer positions with contactless apical domains and intact polarity (and undergo TE differentiation) or are positioned on the inside as surrounded apolar cells that remain pluripotent [8–11]. Following the 16- to 32-cell transition a secondary group of apolar inner cells is similarly supplemented, from outer polarised parental cells, to the primary inner cell progeny population; that together constitute a nascent early blastocyst (E3.5) ICM from which EPI and PrE are derived [12]. Traditionally, relative spatial positioning of blastocyst lineage progenitors was considered via a prism of cell division plane orientation; characterised as those aligning along the embryonic radial axis (i.e. apical-basolateral polarity axis) generating an apolar inner and a polarised outer cell (termed “asymmetric/differentiative” divisions) and those occurring perpendicularly yielding two polarised outer cells (via “symmetric/conservative” cleavages [13]). More recent time-lapse studies indicate most divisions broadly correlate with mitotic spindle alignments along the embryonic radial axis but only a fraction of apolar inner cells are directly deposited post-cytokinesis [14–17]. Indeed, most blastomeres adopt relative spatial positions after oblique-angled divisions, typically resulting in two initially outer residing daughters with significantly unequally sized contactless apical domains. In such situations, increased actomyosin driven cortical tension causes internalisation of cells with smaller contactless apical domains [15, 16]. A role for intra-cellular apical-basolateral polarity in regulating cell internalisation is supported by clonal dysregulation of the apical polarity factor PRKCZ/I resulting in blastomere internalisation [18] and the spontaneous internalisation of naturally occurring apolar outer 16-cell stage blastomeres [15]; also reviewed [19, 20]. Whether relative spatial positioning, and consequent blastocyst cell fate, is essentially a stochastic process subject to intrinsic mitotic spindle angle and cell division plane orientation heterogeneity [21] [22] or subject to other contributing factors (e.g. cell shape, intercellular contact or intrinsic organisation of individual cells [23]) remains unclear. For example, apical to basal repositioning of 8-cell stage nuclei is reported to positively correlate with increased incidence of inner cell generation [24] and positive correlations between polarised contactless apical domain size and mitotic spindle angle orientation along the radial embryonic axis are described [21]; supported by studies on isolated 8-cell blastomeres, that are nevertheless regulated in relative frequency in intact dividing 8-cell stage embryos, [25].

Spatial separation of polarised outer TE progenitors from nascent apolar ICM blastocyst populations defines the “first cell fate decision” [1, 2] and is accompanied by distinct lineage marker gene expression. Outer TE progenitors express the transcription factor CDX2 [26, 27] and early blastocyst ICM cells co-express pluripotency related transcriptional regulators NANOG [28, 29] and SOX2 [11, 30, 31] with the early PrE marker GATA6 [32, 33]; indicative of apparent uncommitted ICM fate preceding the “second cell fate decision”. These expression domains are regulated via differential activation of Hippo-signalling, ultimately supressed in a polarity dependent mechanism in emerging outer TE cells. This is achieved via apical domain sequestration of the Hippo activator AMOT [9, 34] and activated in apolar ICM founders to resist TE differentiation and promote pluripotency [10, 11, 35]. However, early blastocyst ICM is comprised of both primary and secondary ICM founders, each exposed to varying degrees of past Hippo-pathway activation and suppression, thus questioning their relative capacities to preferentially derive EPI or PrE. A debated model contends primary ICM founders, subjected to relatively earlier Hippo-signalling activation, preferentially contribute EPI and secondary inner cells, initially sequestered from active Hippo-signalling and exposed to additional TE-differentiative cues as outer 16-cell stage blastomeres, are strongly biased towards PrE [14, 36–43]. Hence, enhanced mechanistic insight into relative spatial segregation of polarised outer TE-progenitors from both primary and secondary ICM founders will assist understanding of the potentially integrated nature of blastocyst lineage derivation.

The mammalian target of Rapamycin (mTOR) is an evolutionarily conserved serine/threonine kinase of the phosphoinositide 3-kinase family, serving as the central metabolic cellular regulator in response to various intrinsic and extrinsic stimuli. mTOR integrates upstream signalling inputs with downstream effectors, including components of the transcriptional and translational apparatus, to functionally regulate key processes of energy utilisation, specific metabolic pathways, cell-growth and proliferation, autophagy and protein synthesis and degradation. mTOR is the catalytic subunit of two distinct complexes, mTORC1 and mTORC2, respectively regulating cell growth (*e.g.* lipid and nucleotide synthesis, protein synthesis and degradation and autophagy) and survival/proliferation (*e.g.* apoptosis, glucose metabolism, ion transport and cytoskeleton rearrangement). mTOR activity within mTORC1 but not mTORC2 can be inhibited by the compound Rapamycin, whereas second generation ATP analog inhibitors, such as Torin1, are effective against both complexes [44–47]. Active mTORC1 regulates protein translation via phosphorylation of many translation initiation factors and ribosomal related proteins, including the key effectors, eukaryotic translation initiation factor 4E-binding proteins (EIF4EBP1/2/3 [48]). Unphosphorylated EIF4EBPs inhibit 7-methylguanosine-cap (7mG-cap) dependent translation by direct sequestration of the 7mG-cap-binding-complex protein EIF4E from the EIF4F translation initiation complex (also comprising the scaffold protein EIF4G1 and RNA helicase EIF4A [49, 50]). Hence, direct mTOR mediated EIF4BP phosphorylation impairs this inhibitory interaction and facilitates EIF4F initiation complex driven 7mG-cap dependent translation; reviewed [45, 47]. However, sensitivity of specific mRNA translation to mTOR/mTORC1 inhibition (mTORi) is not uniform. Transcripts containing so-called TOP-(5’-UTR terminal oligopyrimidine) or TOP-like sequence motifs (collective referenced here as TOP-motifs), often but not exclusively derived from genes related to protein synthesis itself, are significantly more sensitive to mTORi. Therefore, TOP-motif presence, most conservatively defined as a 7mG-capped C nucleotide followed by a run of 4-15 pyrimidines [51], identifies mRNA transcript classes that are selectively transcribed under conditions of enhanced active mTOR signalling; predominantly via phosphorylation of EIF4BP [49]. Additionally, the mTORC1 substrate and RNA-binding protein LARP1, regulates TOP-motif mRNA translation by directly interacting, in its unphosphorylated state, with TOP-motifs to inhibit translation (an association impaired upon mTORC1 dependent phosphorylation); although this model is disputed (reviewed; [45, 52]).

In the field of mammalian reproduction and early development, mTOR was mechanistically studied during mouse oocyte meiotic development. Pharmacological inhibition of spindle associated mTOR impairs cortical spindle migration, asymmetric extrusion of the first meiotic polar body and causes cytoskeletal disruption [53]. Moreover, mTOR mediated inactivation of EIF4EBP1 (via phosphorylation of specific substrate residues) facilitates appropriate spatio-temporal translation of germinal vesicle enriched or spindle proximal mRNAs important for meiotic maturation, including those with TOP-motifs ([54]; e.g. encoding ANK2 [55]). Post-fertilisation, mTOR directed phosphorylation of EIF4EBP1 is reported as an important translational regulator of the maternal-to-embryonic transition [56]. In early mouse blastocysts (but not earlier cleavage stages), partial pharmacological mTORi (targeting mTORC1 and mTORC2) results in prolonged but reversible and viable *ex vivo* paused development; akin to natural hormonally induced *in vivo* diapause [57]. Additionally, mTOR signalling, operating downstream of active p38-MAPK, is distinctly implicated during PrE (but not EPI) specification from uncommitted early mouse blastocyst ICM populations; whereby defective PrE specification caused by p38-MAPK inhibition [58, 59], is associated with reduced protein translation [60]. Such reports exemplify mTOR associated and lineage specific mechanisms regulating early developmental cell fate, further supported by observations of relative differences in mTOR activity underpinning necessary cellular competition and elimination in early post-implantation embryonic tissues exiting naïve pluripotency [61].

Here we report enhanced levels of M-phase associated phospho-EIF4EBP1 (pEIF4BP1) around the 8- to 16-cell transition, sensitive to mTORi. mTORi around this transition results in the generation of fewer 16-cell stage primary ICM founder cells without affecting apical-basolateral polarity in supernumerary outer cells. Dysregulation of EIF4BP1, LARP1 and EIF4F 7mG-cap-binding-complex function phenocopies mTORi, as does siRNA mediated clonal knockdown of identified TOP-motif containing mRNAs related to the cytoskeleton and secondary RNA structure. These data invoke a mechanism by which mTORC1 activity at the 8- to 16-cell transition facilitates translation of specific TOP-motif containing mRNAs, functionally required to generate primary ICM founder cells; reported to preferentially contribute EPI [14, 38, 40]. However, we also report a lack of a similar active mTOR requirement to generate secondary ICM founders around the 16- to 32-cell transition, suggesting distinct mechanisms of lineage relevant blastomere spatial segregation after successive cleavage rounds. Whilst early blastocysts (E3.5) cultured from the 8-cell stage under mTORi conditions present with fewer ICM cells, resulting late (E4.5) blastocysts (derived when such embryos are transferred to regular culture conditions) recover ICM cell number, specify an appropriately sized EPI but present with evidence of impaired PrE differentiation.

## RESULTS

### mTORC1 signalling during 8-cell stage M-phase entry is associated with generation of primary ICM founders at the 16-cell stage and mTOR-4EBP1-EIF4E/mRNA cap-binding-complex axis function

Motivated by published meiotic phenotypes associated with mTORi during oocyte maturation and defective asymmetric polar body extrusion [53-55, 62, 63], we assayed potential functional mTOR roles during the first spatial cellular separation in preimplantation mouse embryos. We assayed levels of phospho-4EIF4EBP1 (p4EIF4EBP1), a known product of active mTORC1 signalling [64], at the 8- to 16-cell transition and noted increased p4EIF4EBP1 levels associated with 8-cell stage M-phase entry, that were localised around condensing chromosomes and associated with mitotic spindles; returning to basal levels in 16-cell stage progeny (Fig. 1a). Torin1 mediated mTORi showed reduction in pEIF4EBP1 levels, whereas immunofluorescence-staining (IF) for pan-EIF4EBP1 indicated no significant difference in overall EIF4EBP1 expression between control and mTORi treated groups during 8- and 16-cell interphase, although levels did increase after mTORi during M-phase (Figs.1b, S1). We interpret this as indicating an M-phase specific increase in mTOR signalling during the 8- to 16-cell transition. We next determined if this increase was associated with elevated general *de novo* translation, using an O-propargyl-puromycin (OPP) polypeptide incorporation and fluorescent labelling assay. Despite the observed increase in mTORi sensitive pEIF4EBP1 levels, we did not detect a significant difference in *de novo* translation during this period (Fig. 1c); suggesting basal protein synthesis was not affected. We hypothesised observed increases in pEIF4EBP1 levels may not affect global mRNA translation but rather a small subset of functionally significant transcripts, such as those described in oocytes harbouring TOP-motifs [55].

**Figure 1:**
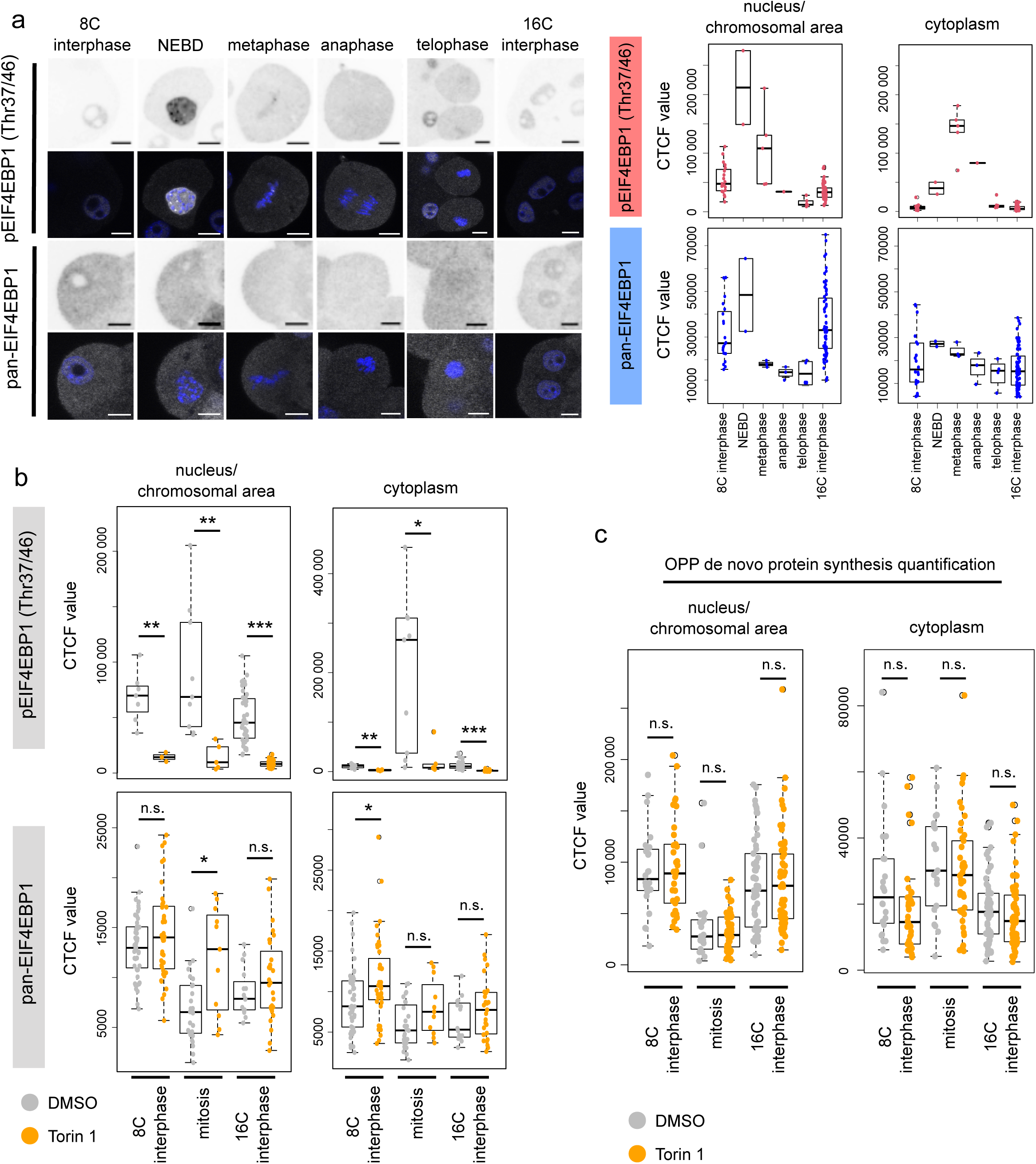
Enhanced mTOR-dependent expression levels of pEIF4BP1 during the 8- to 16-cell cleavage division. **a)** Example IF staining micrographs of pEIF4EBP1 (Thr37/46) and pan-EIF4EBP1 at either 8- or 16-cell interphase and the individual stages of mitotic division from 8- to 16-cell stage in individual blastomeres (left), as quantified separately for nucleus (interphase cells)/chromosomal area defined by DAPI staining (mitotic cells) and cytoplasm (right). EIF4EBP1 visualised in greyscale, DAPI in blue, representative confocal z-stacks for individual stages shown. Scale: 20µm. **b)** Quantification of IF staining of pEIF4EBP1 (Thr37/46) and pan-EIF4EBP1 in 8- and 16-cell interphase and dividing blastomeres, +/-Torin1. **c)** Quantification of nascent translation by O-propargyl-puromycin (OPP) polypeptide incorporation and fluorescent labelling assay in 8- and 16-cell interphase and dividing blastomeres, +/-Torin1.

We noted Torin1 mediated mTORi from the mid-8-cell stage also resulted in significantly fewer inner cells in early-mid- and late-16-cell stage morulae, with multiple examples of embryos lacking any primary ICM founder population, when compared to DMSO treated controls. We also noted no significant differences in the average number of blastomeres that maintained a minimal outer contact, that we termed Small Apical Domain (SAD) cells (Fig. 2a). Furthermore, we observed a similar mTORi phenotype using the alternative inhibitor Rapamycin, suggesting the phenotype is mediated via impaired mTORC1 function ([44–47] – Fig. S2a). Shortening the mTORi exposures (using Torin1) revealed the phenotype of deficient primary ICM founders was minimally centred during a 5 hour window around 8-cell stage mitotic onset and entry into the 16-cell stage (Fig. 2b). Inhibition of the formation of the EIF4F initiation complex using the compound 4EGI1 (that blocks association of the EIF4F mRNA 7mG-cap-binding-complex subunits, EIF4E and EIF4G -[65]) during the same 5 hour window generated a mTORi phenocopy of fewer primary ICM founder cells; confirming the phenotype results from functional dysregulation of the mTOR-EIF4EBP1-EIF4E/mRNA cap-binding-complex axis (Fig. 2b). To further confirm this conclusion, we employed a known dominantly acting recombinant 4EIF4EBP1 construct (in which four known mTOR specific phosphorylation amino acid substrates are mutagenised to Alanine; i.e. 4Ala-EIF4EBP1 – [49]), as its expression should impair elevated levels of 7mG-cap-dependent mRNA translation, irrespective of mTOR signalling status. Accordingly, *in vitro* transcribed mRNA encoding 4Ala-EIF4EBP1 (or wild-type EIF4EBP1 – incorporating a N-terminal HA-epitope tag) was microinjected (plus a lineage injection marker; mRNA encoding Histone-H2B-YFP) into one blastomere of 2-cell stage embryos that were then cultured until the mid-16-cell stage. The expression of the recombinant 4Ala-EIF4EBP1 protein was confirmed by immuno-fluorescent (IF) staining (Fig. S2b). As after mTORi, we observed reduced primary ICM founder cell contribution, restricted to the marked progeny of the microinjected clone, although microinjection of a similar wild-type EIF4EBP1 recombinant mRNA had no significant effect (Fig. 2c). We interpreted these data as further evidence of the involvement of the mTOR-EIF4EBP1-EIF4E/mRNA cap-binding-complex axis as a component of the observed mTORi phenotype. We used a similar microinjection strategy to down-regulate the mRNAs encoding EIF4G1 (one of three subunits of the EIF4F 7mG-cap-binding-complex [64]) and *Larp1* (a mTOR substrate implicated in efficient mRNA translation, particularly of TOP-motif containing mRNAs - [45, 52]) using specific siRNAs (after first confirming efficient siRNA mediated target transcript knockdown by quantitative RT-PCR of siRNA microinjected in 2-cell stage embryos, targeting both blastomeres, cultured to the mid-16-cell stage– Fig. S2c). We again elicited phenocopies previously associated with mTORi involving reduced and clonal primary ICM founder cell contribution (Fig. 2c). Collectively, these data indicate an 8-cell stage and M-phase specific temporal boost in mTORC1 activity, acting via the mTOR-EIF4EBP1-EIF4E/mRNA cap-binding-complex axis, potentiates deposition of daughter blastomeres to the inner compartment of 16-cell stage embryos as primary ICM founding cells.

**Figure 2:**
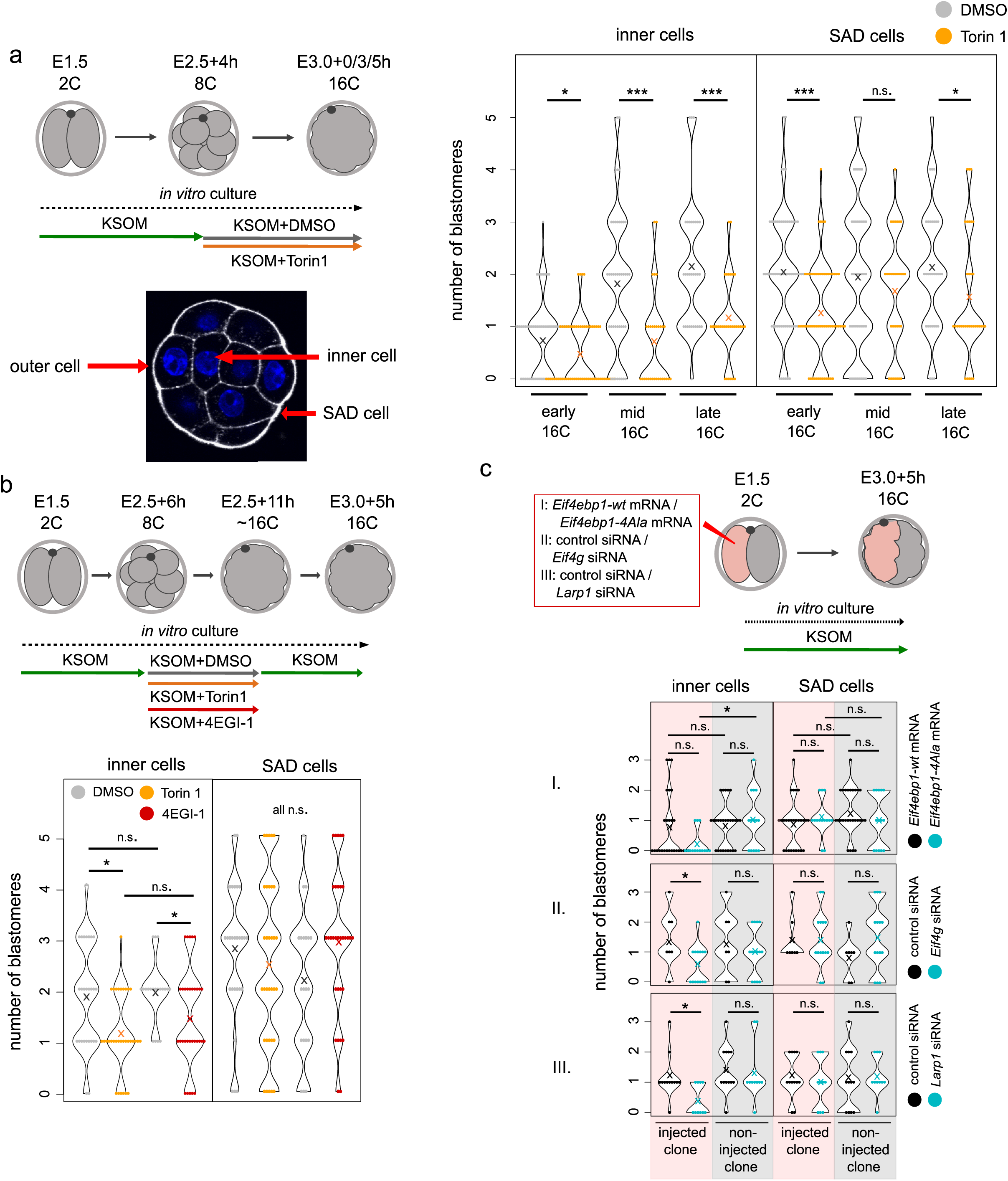
mTOR-regulated 7mG-cap-dependent translation plays a role in primary ICM cell generation. **a)** Experimental scheme, visualisation of example outer, inner and SAD cells, and quantification of the number of inner and SAD cells in 16-cell embryos, +/-Torin1. **b)** Experimental scheme with shorter inhibition period and quantification of inner and SAD cells in 16-cell embryos, +/-Torin1 or +/-4EGI-1. **c)** Experimental scheme and quantification of inner and SAD cells in injected (i.e. indicated siRNA/mRNA) and non-injected clones of 16-cell stage embryos. In all graphs, “x” marks the average value.

### mTORi does not affect apical-basolateral polarisation, relative nuclear positioning upon 8-cell stage M-phase entry nor mitotic spindle orientation

Relative spatial positioning of outer and inner cells from the 16-until the 32-cell stage is known to be tightly regulated by apical-basolateral polarity (established at the late 8-cell stage [13]); whereby in outer blastomeres lacking apical-basolateral polarity, increased relative actomyosin contractility actively segregates cells to the inner compartment [15, 16, 18, 19]. We therefore assayed the expression of polarity related proteins in 16-cell stage embryos exposed to mTORi from the late 8-cell stage using IF; assaying the apical polarity factor aPKC/PRKCZ [18], basolaterally localised cell adhesion protein ECAD/CDH1 [66] and YAP1 (as a readout of polarity dependent Hippo-pathway activity). We did not observe any difference in apical polarity status of embryos with supernumerary outer cells after mTORi and any generated inner cells were appropriately apolar (Fig. S3a). Similarly, ECAD expression was appropriately basolateral in outer cells, although there was a small and significant reduction in the quantified levels of protein expression at such membranes (Fig. S3b). Compared to controls, the relative numbers of blastomeres exhibiting nuclear or cytoplasmic YAP1 expression remained unchanged, indicating correct polarity-dependent regulation of the Hippo-pathway (Fig. S3c). These data indicate the mTORi phenotype of fewer primary ICM founders is not related to regulation of apical-basolateral polarity.

We next employed live fluorescent confocal microscopy embryo imaging to observe individual 8-cell blastomere division. Recovered 2-cell stage embryos were microinjected in both blastomeres with three recombinant mRNAs encoding differentially labelled fluorescent reporter protein constructs (i.e. Histone-H2B-mCherry, GAP43-CFP and alpha-Tubulin-Venus, to visualise chromatin, plasma membranes and tubulin-cytoskeleton/mitotic spindle, respectively – [60, 67]). Microinjected embryos were cultured until the late 8-cell stage and exposed to mTORi (using Torin1) or solvent control DMSO before imaging through the 8- to 16-cell stage transition. We measured relative 8-cell stage nuclei positioning along the embryonic radial axis (determined from the most apical component of each blastomere) upon M-phase entry and the spatial fate of daughter cells by the late 16-cell stage. In control embryos we observed previously reported trends describing increased incidence of inner cell generation associated with more basal nuclear positioning and the generation of two-outer cells being linked with apical positioning [24] (Fig. 3a). In mTORi treated groups, we did not observe any difference in the overall frequency nor distribution of relative nuclear positioning observed in control embryos. However, we found the confirmed trends relating to 16-cell stage blastomere positioning no longer held. Indeed, the observed bias for basal nuclear position yielding an outer and inner/SAD daughter blastomere in the control group is not present in mTORi treated embryos (Fig. 3b). These results indicate previously published outcomes associated with nuclear positioning are nevertheless sensitive to mTORi, even if the frequency and distribution of nuclear positioning was not affected. We next measured relative mitotic spindle angles in relation to the radial axis of each 8-cell stage blastomere (by intersecting a line drawn through the opposing spindle poles and the defined embryonic radial axis - Fig. 3c). Whilst average distributions of relative spindle angles were not significantly different between mTORi and control embryos (Fig. 3d), we observed a significant increase in the generation of two outer daughter cells in mTORi treated embryos arising from acute spindle angles (0-30°) more likely to generate a single outer cell and either a SAD or, more significantly, an inner 16-cell stage blastomere in DMSO treated controls (Fig. 3e). Similarly, we interpret the data to indicate reported mechanisms related to 8-cell stage spindle angle and the propensity for primary ICM founder cell generation [24] are not operative under mTORi conditions, despite not affecting the distribution of observed spindle angles themselves. Moreover, this results in the observed supernumerary populations of outer cells and a deficit of inner cells and strongly implicates a direct role for mTORC1 signalling in post-division positioning of 16-cell stage blastomeres.

**Figure 3:**
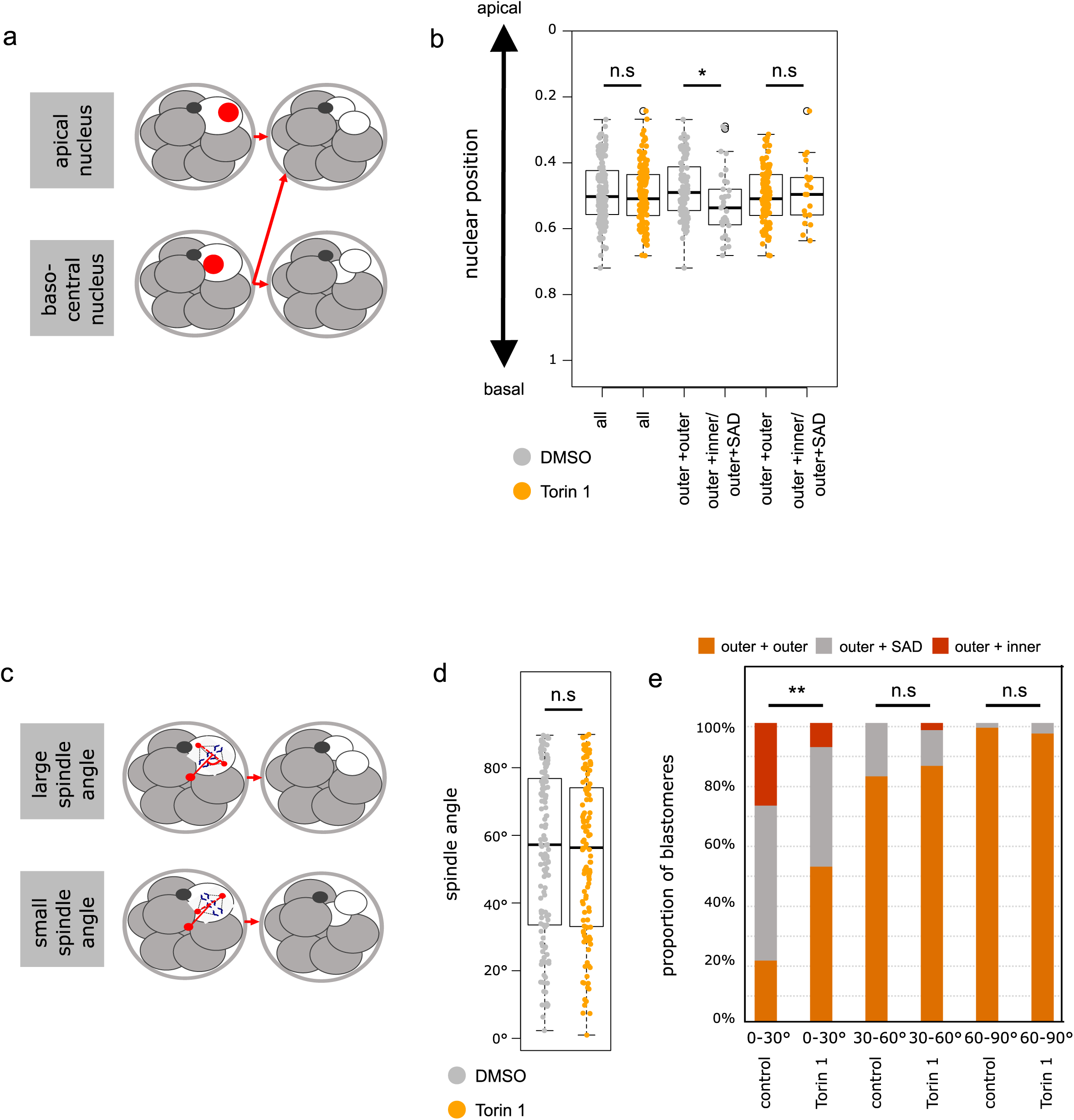
Trends associated with 8-cell stage blastomere nuclear positioning and mitotic spindle angles and the generation of primary ICM cells are not applicable under mTORi conditions. **a)** Scheme of the identified association between 8-cell stage blastomere nuclear position and spatial positions of daughter cells post-division [24]. **b)** Quantification of the position of the immediately pre-M-phase 8-cell stage nucleus on the intra-cellular apico-basolateral axis (i.e. embryonic radial axis), +/-Torin1; in all blastomeres, or those that generated two outer cells or one outer and one inner/SAD daughter cell at the 16-cell stage. **c)** Scheme of the reported association between mitotic spindle angle and spatial positions of daughter blastomeres after 8-cell stage cell division; note, spindle angle is denoted by the angle of lines bisecting the two spindle poles and the radial axis of the embryo from each individual blastomeres most apical membrane domain. **d)** Quantification of the average spindle angle of 8-cell dividing blastomeres, +/-Torin1. **e)** Quantification of the spindle angle versus resulting spatial positions of daughter blastomeres, +/-Torin1. Each bar represents at least 15 blastomeres.

### Dysregulation of candidate TOP-motif containing mRNA generates fewer primary ICM founders

Due to the aforementioned observations (Figs. 1 & 2), we hypothesised that mTOR/mTOC1 might function by regulating translation of TOP-motif containing mRNAs [49, 51, 54]. We identified two TOP-motif containing mRNAs encoding the cytoskeletal proteins; i) Ankyrin-2/ANK2 and ii) Dynactin-2/DCTN2. ANK2 is a protein involved in linking integral membrane proteins to underlying spectrin-actin cytoskeleton [68] that interacts with Dynactin to promote long-range motility of cells [69] and is reported be under regulated translation by mTOR during mouse oocyte meiotic maturation [55]. DCTN2 is a component of the Dynactin macromolecular complex, an interactor of microtubules with reported roles in nuclear positioning and mitotic spindle formation [70] and is also present in a published database of candidate TOP-motif containing mRNAs [49]. After first confirming efficient dsRNA mediated knockdown of the target transcripts at the mid-16-cell stage (Fig. S4 – note, both blastomeres of 2-cell embryos were microinjected with dsRNA and target mRNA levels measured by quantitative RT-PCR), we microinjected one blastomere of 2-cell stage embryos with either *Ank2* or *Dctn2* specific dsRNAs or control dsRNA (targeting GFP), plus an injection marker, and cultured them until the mid-16-cell stage. Morulae were assayed for the contribution of marked clones to primary ICM founder populations. Relating to control GFP dsRNA microinjected embryos, no significant differences of either the marked or unmarked clonal contribution between outer and inner cell populations was observed. Moreover, in *Ank2* and *Dctn2* dsRNA microinjected embryos the non-injected clone did not allocate with any significant difference to equivalent clones in the GFP dsRNA control group. However, primary ICM contribution of the marked *Ank2* and *Dctn2* dsRNA microinjected clone was significantly reduced (Fig. 4). We next employed an empirical mass-spectrometry method to identify further candidate mRNA transcripts, involved in the generation of primary ICM founder cells, by surveying and comparing total detectable proteomes of embryos transiting the 8- to 16-cell stages under control or mTORi conditions (Table S1). Due to the scare amount of sample material, complete proteome coverage was not obtained, but we observed a statistical depletion of DDX21 protein levels (a DEAD box RNA helicase coordinating rRNA transcription/processing during ribosome assembly [71] and a member of a family of helicases implicated in mRNA secondary structure resolution during translation [72]; encoded by an mRNA containing a canonical TOP-motif [49]). Adopting a similar clonal siRNA mediated approach of *Ddx21* transcript knockdown, we again observed a significant reduction in the number of 16-cell stage primary ICM founder cells, that was confined to the marked microinjected clone (Fig. 4) Collectively, the data demonstrate experimental knockdown of candidate TOP-motif containing mRNAs phenocopies mTORi, and direct dysregulation of the mTOR-EIF4EBP1-EIF4E/mRNA cap-binding complex axis, mediated impairment of primary ICM founder cell formation. They also support a hypothesis that enhanced mTORC1 signalling through the 8- to 16-cell transition potentiates translation of specific mRNA transcripts with functional roles in 16-cell stage blastomere spatial positioning.

**Figure 4:**
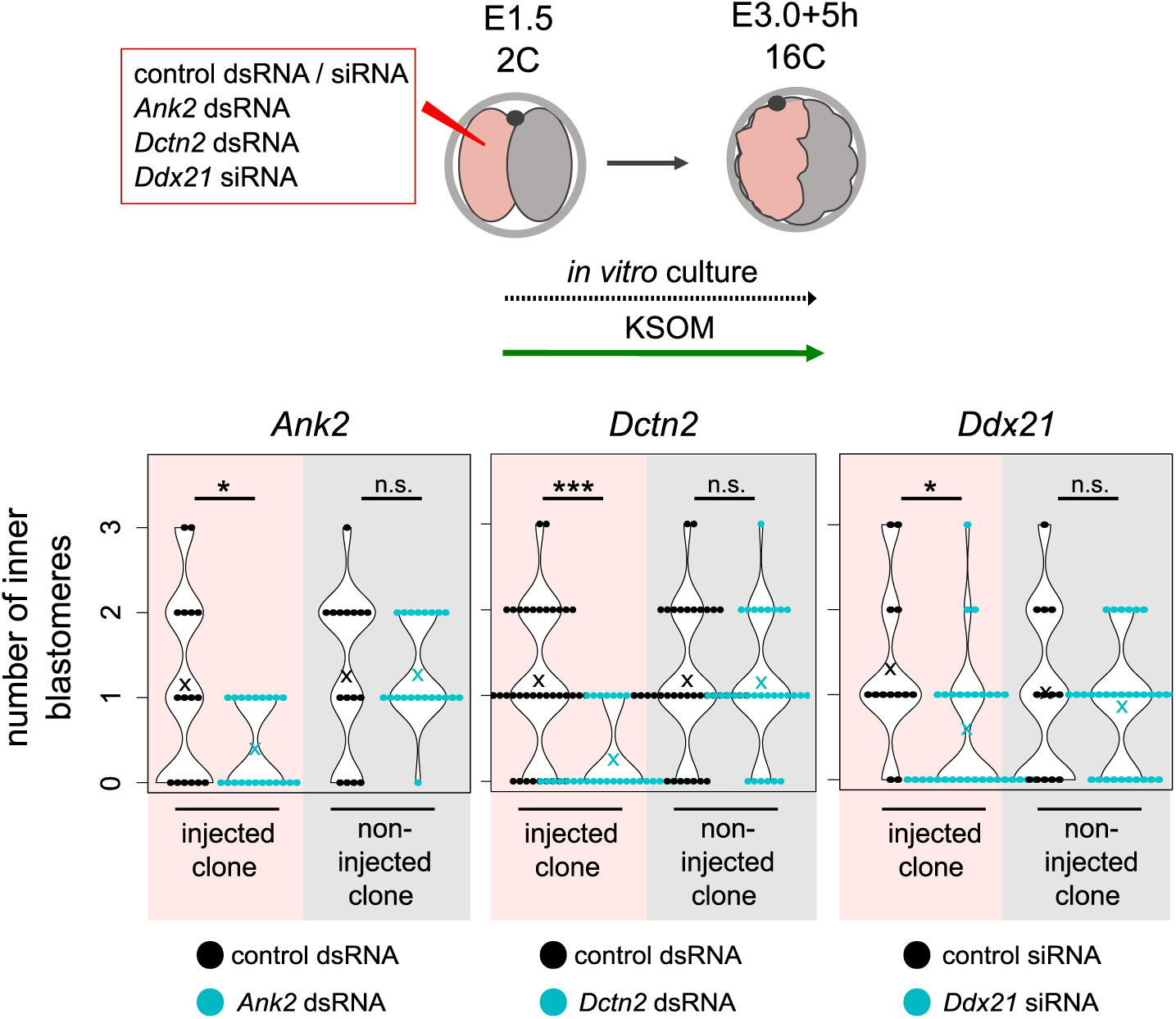
RNAi-mediated knockdown of candidate candidate TOP-motif containing mRNAs also impairs the generation of primary ICM cells. Experimental scheme and quantification of inner and SAD cell numbers in gene/transcript specific RNAi (*Ank2*, *Dctn2* and *Ddx21*) injected and non-injected clones in 16-cell stage embryos. In all graphs, “x” marks an average value.

### mTORi impairs 16-cell stage primary but not 32-cell stage secondary ICM founder cell generation

We next asked the question whether mTORC1 dependency during primary ICM founder cell generation was applicable to generation of secondary founders, after the 16- to 32-cell transition. Embryos were exposed from the pre-M-phase late-8-cell stage to mTORi (with Torin1) or solvent control (DMSO) and cultured to the mid-32-cell stage (defined morphologically as control blastocysts exhibiting cavities occupying ∼50% of embryo volume). mTORi treated groups displayed significantly fewer inner/ICM cells than controls but to a lesser extent (i.e. as could solely be explained by the initial lack of dividing 16-cell stage primary founder ICM cells); suggesting mTORi did not impair internalisation of secondary ICM founder cells (Fig. 5a). Indeed, if mTORi/DMSO was administered from the mid-16-cell stage, no significant differences in ICM cell number were observed (Fig. 5a), indicating a developmentally staged requirement for enhanced mTORC1 activity to appropriately segregate primary ICM founder cells that is not required during the subsequent 16- to 32-cell transition. When mTORi was limited to a period spanning the late-8-cell stage until the mid-16-cell stage and embryos further cultured under normal conditions to the mid-32-cell stage, the total number of primary inner cells (as mathematically calculated) was still impaired although the total number of inner cells was now restored to levels statistically insignificant to DMSO treated controls (although still slightly fewer), indicative of a degree of compensatory secondary ICM founder cell generation post-Torin1 removal (Fig. 5a). We next asked if mTORi would affect the specification of peri-implantation stage (E4.5) blastocyst lineages, as literature reports have suggested primary ICM founders are biased to form EPI and secondary ICM founders PrE [14, 36, 38, 42, 43]. Pre-M-phase late 8-cell stage embryos were exposed to either mTORi (using Torin1) or DMSO control culture conditions until the mid-32-cell stage (concomitant with irreversible TE specification [73]) and then transferred to conventional culture media until the late blastocyst stage (E4.5); note, mTORi could not be given beyond this point as developmental diapause would result [57]. Blastocysts were then IF stained for CDX2, GATA4 and NANOG, as markers of specified TE, PrE and EPI lineages, respectively. We observed the ICM cell number was equivalent between the two groups but overall cell number in mTORi treated embryos was less, possibly indicating reduced cellular fitness in the TE cell lineage. Within the ICM, numbers of NANOG+ and GATA4-cells (indicative of EPI specification) were equivalent under mTORi versus control conditions but the number of NANOG- and GATA4+ cells (indicative of PrE differentiation) was reduced, but not significantly. Although, we also observed a significant increase in the percentage of ICM atypically co-expressing both markers (i.e. NANOG+ and GATA4+; Fig. 5b), also indicative of perturbed PrE differentiation. We speculated if recovered ICM cell number in mTORi late embryos, that present with fewer ICM cells at the mid-32-cell stage, may result from reduced levels of confirmed ICM apoptosis known to occur during blastocyst maturation [38] but IF staining for cleaved Caspase-3 (at E4.0 +7 hours), did not reveal any significant decrease in mTORi treated embryos compared to DMSO controls (Fig. 5c). These data indicate mTORi during the 8- to 32-cell stages, including confirmed deficits in primary ICM founders, is compensated during blastocyst maturation to ensure an appropriately sized ICM consisting of correctly specified EPI but impaired PrE differentiation. They also suggest reduced numbers of primary ICM founders do not ultimately impair EPI specification that may be inferred from other reports linking biased EPI and PrE formation to primary and secondary ICM founders, respectively [38, 40]; although the extent to which regulatory compensatory mechanisms, not ordinarily operative in unperturbed development, participate cannot be excluded.

**Figure 5:**
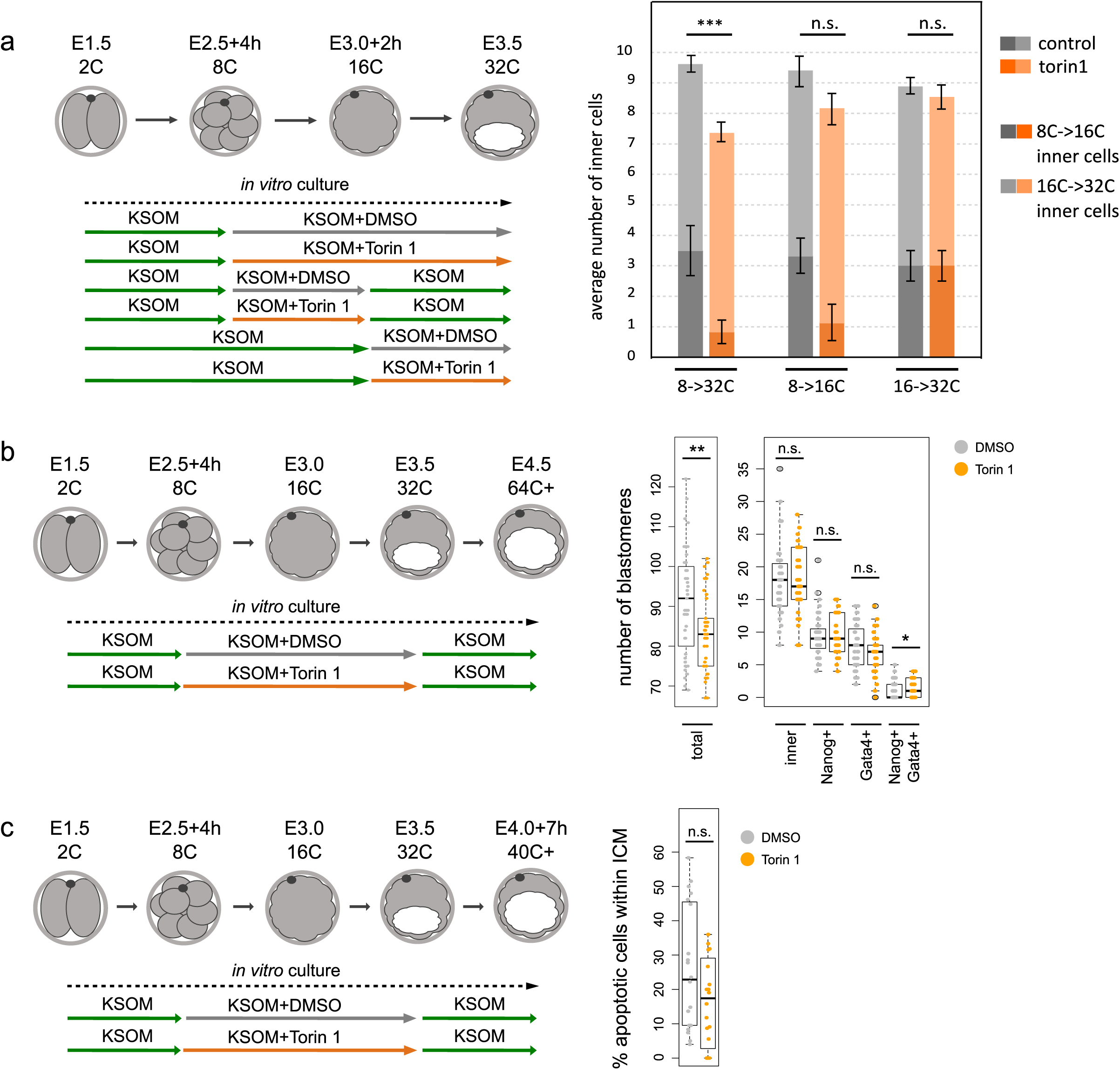
mTORi does not affect secondary ICM founder cell generation and ICM cell numbers recover during blastocyst maturation (E3.5-E4.5) after prior mTORi from the late 8-cell stage. **a)** Experimental scheme and quantification of inner cell numbers at 32-cell stage, as contributed by primary and secondary ICM cells, +/-Torin1. Primary ICM cell count was estimated by fixation of some embryos at 16-cell stage and quantification of their number of inner cells, to allow for the number of such primary and secondary founders to be determined at the 32-cell stage, when deducting this number and allowing for the extra cell division. Each condition represents at least 25 embryos. **b)** Experimental scheme and quantification of total cell number, ICM cell number and the numbers of NANOG+ GATA4-(EPI), NANOG-GATA4+ (PrE) and NANOG+ GATA4+ ICM cells, +/-Torin1 (E4.5). **c)** Experimental scheme and quantification of the proportion of apoptotic ICM cells (positive for cleaved Caspase-3 IF staining), +/-Torin1 (E4.0+7h).

### Supernumerary outer cells in mTORi treated blastocysts exhibit ICM-like marker gene expression

We noted embryos exposed to mTORi (using Torin1) from the pre-M-phase late-8-to the 32-cell stage did not exhibit any significant difference in average outer cell numbers appropriately expressing CDX2 or nuclear sequestered YAP1 (indicative of suppressed Hippo-signalling), despite having supernumerary outer cells (and fewer ICM cells); Figs. 6a, b & S5a. We speculated this might reflect differences in contactless apical domain area. We measured the length of apical domains (in maximal confocal z-sections) in all outer cells and found atypical CDX2-blastomeres in the mTORi treated group have significantly smaller apical domains than CDX2+ cells. Moreover, that this average apical domain length was indistinguishable from that in spontaneously occurring CDX2-blastomeres in DMSO controls (Fig. 6c), albeit occurring in more outer blastomeres (a small and significant reduction in apical domain size in CDX2+ cells in the mTORi group versus the DMSO group was also observed, possibly reflecting mTORi embryos having supernumerary outer cells). We next categorised subcellular localisation of YAP1 in outer cells (as either; i. exclusively nuclear, ii. cytoplasmic and nuclear or iii. only cytoplasmic – as a readout of Hippo-signalling activity) and compared average apical domain size in DMSO and mTORi treated embryos (Fig. 6c). No significant difference in outer blastomeres with appropriately exclusively nuclear YAP1 (associated with suppressed Hippo-signalling and TE differentiation) was seen. Neither was there any difference between the two groups in outer blastomeres with only ectopic cytoplasmic YAP1 localisation. However, the average apical domain size of such cells was robustly and significantly smaller when compared to those with exclusively nuclear YAP1 within each group (again more frequently observed in the mTORi group). We term these cells MAD (Medium Apical Domain), to distinguish them from SAD cells with only minimal contactless apical domains and interpret the data reflecting a threshold in apical domain size needed to supress Hippo-signalling. Consistently, we also observed significant, yet intermediary, reductions in apical domain size in mTORi treated embryos correlating with YAP1 localisation in both the cytoplasm and nucleus (not observed in DMSO controls); again, possibly relating to such embryos having supernumerary outer cells after mTORi. These data indicate smaller apical domain sizes of MAD cells, either occurring infrequently and spontaneously in DMSO controls or with increased incidence under mTORi conditions, correlates with increased Hippo-signalling (i.e. cytoplasmic YAP1) and a lack of CDX2 expression/TE differentiation. We next assayed, in each group, extents of apical domain polarity via quantitative IF (normalised to measured apical domain area) against the polarity factor PARD6B [4], as a function of CDX2 expression. In control embryos there were no differences in apical polarity between infrequently observed CDX2-MAD and appropriately CDX2+ outer cells. However, after mTORi we found supernumerary CDX2-outer MAD cells either exhibited apical polarity to the same extent as control group outer cells (irrespective of CDX2 status) or it was actually increased (averaging an overall significantly higher level than in CDX2+ cells of the same embryos; Fig S5b). Indeed, when PARD6 expression was measured on the whole embryo level, there were no significant differences between DMSO or mTORi treated embryos (Fig. S5c). These data indicate the failure of supernumerary outer MAD cells to activate CDX2 expression (and suppress Hippo-signalling) after mTORi is not associated with defective apical polarisation at the 32-cell stage and that such cells cannot appropriately specify TE in a manner germane to relative spatial position or polarity status. We hypothesised the MAD cell phenotype was caused by insufficiently large, albeit still polarised, apical domains being unable to appropriately sequester and functionally inhibit the essential embryo Hippo-pathway activator AMOT from basolateral membrane signalling domains [9, 34]; thus, conferring an ectopic pseudo-inner cell phenotype, that due to the presence of apical-basolateral polarisation prevents blastomere internalisation. Using embryos exposed to mTORi or DMSO control from the pre-M-phase late-8-to the 32-cell stages, we assayed for ectopic localisation of AMOT on outer cell lateral membranes of CDX2+ and CDX2-outer cells (note, microscopic resolution prevented objective assessment of basal AMOT localisation due to proximity of inner cells with normal plasma membrane associated AMOT). However, in both groups we did not detect enhanced basal AMOT localisation in mTORi treated embryos versus DMSO controls and neither in MAD cells (Fig. S5d).

**Figure 6:**
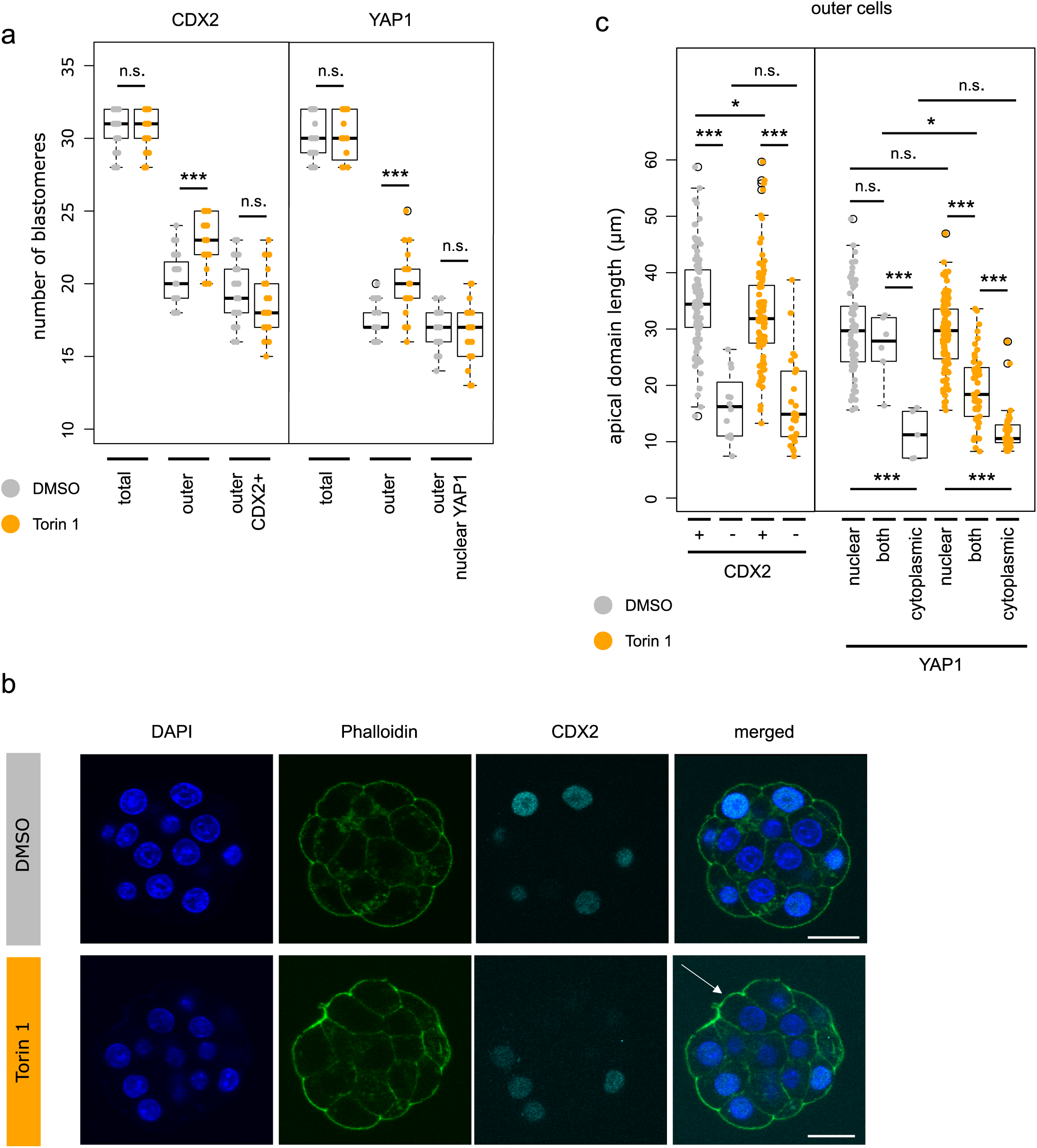
Supernumerary outer cells in mTORi 32-cell stage embryos exhibit molecular characteristics of ICM-like cells. **a)** Quantification of total number of blastomeres, outer blastomeres, and CDX2 + outer blastomeres, or outer blastomeres with nuclear YAP1 without cytoplasmic YAP1 signal, +/-Torin1. **b)** Example IF staining micrographs of CDX2 staining in 32-cell stage embryos cultured under control and mTORi conditions. CDX2 visualised in cyan, DAPI in blue, phalloidin in green, representative confocal z-stacks for individual stages shown. Scale: 20µm. An outer cell without CDX2 in mTORi condition marked by an arrow. **c)** Quantification of apical domain length in single z-stack (with largest membrane length) in outer cells +/-CDX2 and nuclear, cytoplasmic or nuclear + cytoplasmic YAP1, +/-Torin1.

Collectively, these data indicate mTORi induced supernumerary MAD outer cells are unable to inhibit Hippo-pathway activation that would ordinarily permit nuclear accumulation of YAP1 and TE differentiation. However, whilst such blastomeres display smaller contactless apical domains, they nevertheless remain polarised and capable of sequestering AMOT. Therefore, the mechanism by which MAD outer cell activation of Hippo-signalling occurs and results in a pseudo-inner cell phenotype must be independent of apical-basolateral polarity and AMOT itself; potentially related to increased neighbouring MAD cell contacts. However, the fate of such MAD cells and their progeny by the late peri-implantation blastocyst stage (E4.5) remains to be resolved.

## Conclusions

We confirm enhanced mTORC1 signalling levels around the onset of M-phase in 8-cell stage mouse embryo blastomeres, that via regulation of the mTOR-EIF4EBP1-EIF4E/mRNA cap-binding-complex axis, positively influences specific generation of primary, and not secondary, populations of ICM founder cells. Although consistent with published reports associated with regulation of apical basolateral polarity, pre-M-phase positioning of 8-cell stage nuclei and resulting mitotic spindle orientation affecting primary ICM founder generation [15, 18, 21, 24, 25], we do not observe any directly consistent and explanatory perturbations in these processes. Rather, we propose mTORi mediated mis-localisation of 16-cell stage blastomeres is not directly related to such mechanisms *per se*, but manifest in impaired translation of functionally significant and specific subsets of mRNA, normally intransigent to protein translation under basal mTORC1 signalling (including, but not necessarily limited to, those containing TOP-motifs). The fact embryos partially compensate their development during blastocyst maturation (ensuring appropriate EPI specification but impaired PrE differentiation) after prior mTORi during the pre-M-phase 8-to mid-32-cell stages, is testament to their known developmental regulative capacity and maybe linked to ectopically activated Hippo-signalling in supernumerary 32-cell stage MAD cells (unrelated to the classically recognised apical-basolateral polarity model of sequestered AMOT localisation [9–11, 34]).

## DISCUSSION

Considerable debate exists regarding developmental history of ICM cells and their eventual fate as specified EPI or differentiating PrE (i.e. derivation of primary and secondary ICM founders). Non-invasive timelapse and lineage tracing analyses of transgenic reporter embryos have suggested primary ICM founders are biased to form EPI (in 75% of cases) and secondary ICM cells are strongly fated to yield PrE (in 85% of cases); with infrequent third-wave ICM allocation invariably fated to PrE [38]. Thus, supporting a developmental history model of ICM fate (also reviewed; [36, 37]). This is contested by similar independent lineage analyses, permanently marking cell clones, revealing a lack of developmental history effect on EPI and PrE derived post-implantation tissues [39]. Subsequent comparative reanalyses of these datasets have highlighted a context dependency regarding initial numbers of primary ICM founder cells. For example, the former study reported an average of 2.8 generated primary ICM cells (contributing ∼50% of the eventual 32-cell stage ICM), whilst the latter described formation of 4.8 such cells (resulting in 80% contribution to 32-cell stage ICM [14]; an increase possibly related to inadvertent microinjection mediated down-regulation of 8-cell stage apical-basolateral polarity, known to promote cellular internalisation via enhanced actomyosin contractility [16, 18]). Such reappraised data raise the possibility that when 16-cell stage primary ICM founder cells are limiting, progeny are biased to form EPI but surplus contribution can spill over into PrE generation. Consistently, plots of relative percentage contributions of generated primary ICM cells to total ICM number and their eventual EPI or PrE fate support this model [14, 39]. Hence, developmental history and the primary/secondary origin of ICM founder cells seems able to affect ultimate EPI versus PrE fate but is subject to context dependent regulative parameters. This is further supported by molecular observations reporting elevated levels of *Fgf4* and *Fgfr2* transcripts in primary and secondary ICM founders, respectively [31, 42]. We report elevated mTOR signalling upon 8-cell stage mitotic entry facilitates primary ICM founder generation but importantly is not required during formation of secondary ICM populations. Restriction of mTORi sensitivity to primary ICM cell formation is significant, implying distinct mechanisms underpinning generation of successive ICM founder cell populations that may be consequential to their eventual fate under unperturbed development. However, surprisingly when embryos were cultured under mTORi conditions from the pre-M-phase late-8-to 32-cell stages, followed by blastocyst maturation in conventional media, we found the recorded deficits in 32-cell stage ICM cell numbers (solely originating from reduced primary ICM founder contribution) were not only compensated to levels seen in control embryos (although the TE cell number was reduced) but the overall number of ICM cells correctly specified as EPI (i.e. NANOG+ and GATA4-cells) was also statistically the same; although PrE differentiation was impaired (i.e. combined reduced NANOG-/GATA4+ and enhanced NANOG+/GATA4+ cell numbers); Fig. 5. Such data may seem counterintuitive regarding the developmental history model, as it might be expected fewer primary ICM founders would result in less EPI cells. However, it may also simply reflect developmental regulatory capacity ensuring appropriate pluripotent EPI formation (as a foetal progenitor pool) at the expense of PrE differentiation; a mechanism that during unperturbed development is not normally executed. This is possibly related to lower levels of expressed FGF4, required to drive PrE differentiation [39], in the smaller ICMs of mTORi treated early blastocysts. It is therefore impossible to unequivocally conclude if 8-to 16-cell active mTOR dependent mechanisms of facilitating primary ICM founder generation supports such a developmental history related model. Although previously, we reported clonal inhibition of TE cell fate, using microinjected siRNAs specific for *Tead4* transcripts, generates excess ICM contribution favouring EPI and biased against PrE formation, respectively [43]; strongly suggesting an extra ∼12 hours of polarity-dependent Hippo-pathway suppression in outer 16-cell stage blastomeres may prime derived secondary ICM founders to preferentially differentiate towards PrE. Additionally, a recent paper reports NANOG dependent coordinated expression of pluripotency related gene expression during the 16-to 32-cell stage transition, suggesting EPI specification may actually originate (in part) in primary ICM founders [74]. Whilst not definitive, these collective data at least support an aspect of developmental history underlying ICM cell fate, that is nonetheless potentially subject to regulative and compensatory mechanisms. In this context, it is possible individual dividing 8-cell blastomere mTOR activity may act as a developmental rheostat (potentially responding to cellular metabolic status) to regulate derived primary ICM founder numbers, that can then be fine-tuned by distinct and presently unknown and/or stochastic mechanisms (e.g. involving of cell-cell contacts). Thus providing embryos two opportunities to generate a germane number of ICM progenitors necessary to successfully sustain both EPI and PrE specification.

Another aspect of developmental regulation arising from mTORi from the pre-M-phase late-8-to the 32-cell stages, are observations of supernumerary MAD outer cells that fail to express CDX2 and exhibit atypical cytoplasmic localisation of YAP1, indicative of pseudo-inner cell Hippo-pathway activation and failed TE differentiation, despite exhibiting intact apical-basolateral polarity; strongly suggesting MAD cells are undergoing regulative adaptation. As such CDX2-MAD cells exhibit statistically smaller apical domains, we considered this may result in an insufficient capacity to sequester the Hippo-pathway activator AMOT, despite their polarised status. However, we could find no evidence of ectopic lateral membrane AMOT localisation that would support this model (Fig. S5). Nevertheless, the fact mTORi induced formation of supernumerary outer MAD cells is associated with failed TE specification and activated Hippo-signalling (Fig. 6) indicates existence of additional mechanisms to activate Hippo-signalling independently of classically described polarity dependent functional sequestration of AMOT to the apical domain [9–11, 34]. Moreover, the fact MAD cells have reduced contactless apical domains suggests these are potentially related to mechanisms reliant on enhanced cell contact or mechanical force akin to other described contexts of Hippo-signalling regulation; e.g. regulation of tissue growth and organ formation (reviewed [75]). Although we did observe small yet significant reductions in the basolaterally localised cell adhesion protein CDH1 in 16-cell stage embryos after mTORi (Fig. S3), it will be interesting to investigate to what extent described molecular factors underpinning these polarisation independent mechanisms affect relative spatial blastomere positioning during unperturbed and regulative preimplantation mouse embryo development.

In embryos cultured under mTORi conditions from the pre-M-phase late-8-until the 32-cell stages and then cultured in conventional media to the late blastocyst (E4.5) stage, we observed impaired PrE differentiation manifest as reduced NANOG-/GATA4+ and enhanced NANOG+/GATA4+ ICM cell numbers (although the number of specified EPI cells, compared to control groups, was equivalent; Fig. 5). The presence of GATA4+/NANOG+ cells, ordinarily are not observed in control embryos, is notable and possibly resembles an uncommitted cell fate state reminiscent of NANOG and GATA6 co-expression in early blastocyst ICM [76], or an attempt to initiate PrE differentiation without downregulating pluripotency. Interestingly, we previously reported such atypical NANOG and GATA4 co-expression during mouse blastocyst maturation under p38-MAPK inhibited (p38-MAPKi) culture conditions [58–60]; particularly when p38-MAPKi is supplemented by pharmacological activation of mTOR [60]. Such phenotypes are associated with impaired general protein synthesis and associated reductions in polysome formation and rRNA processing [60]. Consistently, we find if mTORi from the pre-M-phase late-8-cell stage is substituted with p38-MAPKi, we can elicit the same phenotype of fewer primary ICM founder cells (Fig. S5e); further cementing an emerging importance of p38-MAPK function and its link to wider mTOR signalling in preimplantation development. However, the exact mechanistic details require further investigation.

The mTORi phenotypes of impaired primary ICM founder cell formation described here are mechanistically novel; neither being related to apical-basolateral polarity defects nor positioning of pre-M-phase 8-cell stage nuclei along the embryonic axis or altered orientation of mitotic spindles ([24]; Figs. 3 & S3)) and indicate functionally downstream mTORC1 mediated mechanisms regulating relative blastomere spatial positioning at the onset of the 16-cells stage. Our data denote mTORi phenotypes are, at least partially, based on temporally controlled mTORC1 regulated translation of subsets of functionally significant mRNAs, involving the mTOR substrates EIF4EBP1, LARP1 and assembly of the 7mG-cap-binding-complex (EIF4F). Moreover, identification and functional verification of candidate TOP-motif containing transcripts encoding ANK2, DCTN2 and DDX21 (eliciting similar primary ICM founder cell deficits, after clonal RNAi mediated knockdown; Fig. 4) indicates mTORC1 can regulate relative 16-cell stage blastomere positioning via potentiating specific TOP-motif containing mRNA translation (although other classes of mRNA may be affected); but exclusively at the 8-to 16-cell stage transition. As ANK2 and DCTN2 are cytoskeletal proteins [68, 70] such mechanisms likely involve cytoskeleton remodelling. Moreover, identification and validation of DDX21, an RNA binding helicase implicated in rRNA processing and potential resolution of RNA secondary structure [71, 72], is also consistent, as most characterised TOP-motif containing mRNAs encode proteins related to protein synthesis itself (reviewed: [77]). Thus, suggesting the possibility DDX21 levels contribute a positive feedback loop ensuring efficient translation of other, potentially TOP-motif containing, mRNAs. The phenocopy of mTORi mediated deficits in primary ICM cell formation using p38-MAPKi (Fig. S5e) is also notable, as we previously identified DDX21 as a p38-MAPK effector protein in early mouse blastocysts development and a verified component of PrE specification [78]. Enhanced mTOR mediated translation of *Ank2* transcripts, in proximity to condensing chromosomes and forming meiotic spindles, is reported in maturing mouse oocytes and ensures appropriate and highly asymmetric cell divisions generating the first polar body [54, 55]; implying potential mechanistic similarities that generate primary, but not secondary, populations of blastocyst ICM founders. The molecular signal tightly regulating functionally elevated levels of mTORC1 activity at the onset of 8-cell stage M-phase currently remains unknown. An obvious candidate is the cell-cycle dependent kinase CDK1. Increased CDK1 activity upon mouse oocyte meiotic resumption is proposed as a potential trigger for increased mTOR activity [62], possibly involving PLK1 [79]. However, functional verification of this hypothesis in cleavage stage embryos is hindered as pharmacological inhibition of either kinase results in M-phase arrest. Notwithstanding, our data collectively illustrate temporal regulation of mTORC1 activity, and the translational control of specific and functionally significant mRNA transcripts, as an emerging theme during preimplantation stage mouse embryo development. It will be of great interest to develop these findings, for example utilising emerging contemporary techniques optimised to directly assay single cell/embryo translatomes, reflecting mRNAs under-going direct translation, under control and mTORi or p38-MAPKi culture conditions (e.g. using Scare Sample Polysome profiling – SSP [80], or Ribo-ITP (Tono et al., 2021) combined with mRNA-Seq) and in other mammalian preimplantation embryo species.

## MATERIALS & METHODS

### Superovulation and embryo isolation

All animal work was conducted in accordance with Act No 246/1992 Coll., on the protection of animals against cruelty under the supervision of the Central Commission for Animal Welfare, approval ID 51/2015. Derivation of all experimental embryos was conducted according to the following protocol, unless specifically stated otherwise. As previously described [43], F1 generation of 8-week female hybrid mice (generated via C57BL6 female and CBA/W strain crosses) were intraperitoneally injected with 7.5IU of PMSG (pregnant mare serum gonadotrophin; Merck) and reinjected after 48 hours 7.5IU hCG (human chorionic gonadotrophic hormone; Merck), before overnight mating with F1 or F1 hybrid transgenic mT/mG stud males (original mT/mG transgenic mouse line obtained from Jackson Laboratories, STOCK Gt(ROSA)26Sortm4(ACTB-tdTomato,-EGFP)Luo/J., https://www.jax.org/strain/007576 -[81]; expressing a membrane associated Tomato fluorescent reporter, that when exposed to Cre-recombinase would be excised and replaced by a membrane GFP reporter – in the case of AMOT IF experiments). At least 4 hours before dissection, culture dishes were prepared with twenty 10μl drops of KSOM+AA medium (Embryo-Max; Millipore), covered with mineral oil (Irvine Scientific) and equilibrated in an incubator at 37°C and 5% CO_2_ atmosphere. 2-cell stage (E1.5) embryos were recovered (45-47 hours post-hCG) into and washed through, on a heated stage, 20μL drops of prewarmed (37°C) M2 media containing 4mg/ml BSA (bovine serum albumin - Merck) and then transferred through the series of KSOM+AA culture drops of pre-equilibrated plates, before transfer into the incubator (37°C and 5% CO_2_) and *in vitro* culture to the desired developmental stage. In the case of time-lapse confocal microscopy live embryo imaging, 2-cell stage embryos were similarly recovered from hybrid superovulated females (using 5IU of PMSG and 5IU of hCG; Merck), generated after crossing BDF1 male and CD1 females (each strain obtained from Anlab, Czech Republic), prior to microinjection of recombinant mRNAs (see below). Embryos were then cultured as described in KSOM+AA.

### Embryo inhibitor treatments

In each experiment, embryos at the desired stage were divided into two equal groups (one for vehicle controls and the inhibitor treatment group). 4EGI-1 (Merck), Rapamycin (Merck) and Torin1 (Selleckchem), diluted in DMSO, were used at final KSOM+AA culture concentrations of 100μM, 5μM and 20μM, respectively (with vehicle controls consisting of an equivalent volume of supplemented DMSO). Embryos were then *in vitro* cultured to the desired assay point +/-inhibitor or transferred into non-supplemented conventional KSOM+AA media for further culture, before being processed for the appropriate assay. All inhibitor/vehicle control KSOM+AA culture plates were pre-equilibrated at 37°C in a 5% CO2 atmosphere for at least 4 hours prior to addition of embryos.

### 2-cell stage embryo microinjection

Preimplantation mouse embryo microinjection was performed as previously described [43]. Specific predesigned Silencer-Select gene siRNAs (ThermoFisher Scientific; *Eif4g* - s101902*, Larp1* - s91534, *Ddx21* - s80158 or All-Stars negative murine control non-targeting control/NTC, from Qiagen were microinjected at a final concentration of 10µM each). dsRNAs (targeting *Ank2*, *Dctn2* or GFP mRNAs) were in-house synthesised from T7 RNA polymerase promoter-linked PCR products, using the MEGAscript T7 *in vitro* transcription kit from ThermoFisher Scientific according to provided instructions, and were microinjected at a final concentration of 200ng/μl. Templates for dsRNAs generation were generated using the following PCR oligo primer-pairs: *Ank2* sense – taatacgactcactatagggCCTCATCGAATGCCTCACCA, anti-sense – taatacgactcactatagggTTCTCCTTGGCAGCACAGAG and *Dctn2* sense – taatacgactcactatagggGGCATTGCCAGGAATGAG, anti-sense – taatacgactcactatagggCTGTCCTCTTGGTCTTTCCAA, as designed by ERNAi design tool [82] and GFP sense taatacgactcactatagggAGAGTACAAATTTTCTGTCAGTGGAGAGG, anti-sense taatacgactcactatagggAGATGTATAGTTCATCCATGCCATGTGTA).Recombinant mRNAs were generated from T3 mediated *in vitro* transcription/IVT of cDNA inserts cloned into the vector pRN3P [83], incorporating 5’ and 3’ UTRs from the frog beta-globin gene for enhanced stability, using the ThermoFisher Scientific mMESSAGE mMACHINE T3 and poly-A-tailing kits, as instructed. mRNAs were microinjected at following concentrations: encoding HA-4Ala-EIF4BP1 200ng/µl (inert derived from [49]) or wild-type HA-EIF4BP1 200ng/µl (cloned in-house) and Histone-H2B-mCherry/YFP 50 ng/µl (cloned in-house) - as fluorescent reporter genes/confirmed microinjection markers [84]). RNAs were microinjected in either single, or both, 2-cell stage embryo blastomeres (to generate control or gene specific dysregulated clones representing 50% of embryonic cells, or the entire embryo for Q-RTPCR confirmed assessment of RNAi mediated target gene knockdown, respectively) in suspended M2 + BSA media drops, overlaid with mineral oil using IX71 inverted-microscope (Olympus), micromanipulators (Leica) and FemtoJet microinjection system (Eppendorf). As a microinjection/clonal lineage tracer marker, all siRNA/mRNAs were co-injected with either Rhodamine-conjugated dextran beads (RDBs; final concentration 2 μg/μl) or Histone H2B-RFP/YFP encoding recombinant mRNA (derived from IVT of cloned cDNAs in to pRN3P, as described above [83]) Non-microinjected embryos (1–3 per experiment) served as embryo culture sentinels for subsequent appropriate *in vitro* development in KSOM+AA (as described above). Regarding time-lapse confocal microscopy live embryo imaging experiments, recovered 2-cell stage embryos were microinjected in both blastomeres, using a IM-300 Narishige microinjector on a Leica DM IL inverted microscope, with the following recombinant mRNAs encoding fluorescent reporters (prepared by IVT of pRN3P plasmid templates as described above): Histone-H2B-Venus, GAP43-CFP and alpha-Tubulin-Venus, each microinjected at 10 ng/µl. Note, the correct size and integrity of IVT generated mRNA/dsRNA constructs were first confirmed on denaturing and regular agarose gels, respectively, prior to microinjection.

### Embryo fixation and immuno-fluorescent and fluorescent phalloidin staining

Protocols were as previously described [43]; briefly, prior to fixation embryonic *zona pellucidae* were removed in prewarmed (37°C) drops of Acid Tyrode’s solution (Merck) diluted in M2. Embryos were then fixed, on 1.5% agar coated culture dishes, in 20μl drops of a 4% para-formaldehyde solution (PFA; Santa Cruz Biotechnology), overlaid with mineral oil, for 20 minutes at 37°C. All subsequent steps were conducted at room temperature, unless otherwise stated. In 96-well micro-titre plates, embryos were then washed through three 70 μl drops of PBST (phosphate-buffered saline with 0.15% Tween 20 - Merck) and placed in 50 μl of 0.5% Triton-X100 (Merck) permeablisation solution diluted in PBS for 20 minutes. Embryos were then washed through three 70μl drops of PBST before being transferred to 50μl drops of blocking 3% BSA (Merck) in PBST for 30 minutes. Desired primary antibody dilutions (see below) were prepared in 3% BSA PBS-T (in 5μL volumes) solution into which embryos were transferred for overnight incubation at 4°C (overlaid with mineral oil). Following three washes through 70μL drops of PBST, embryos were subject to a secondary 3% BSA block (1 hour) and then transferred into 5 μl 3% BSA drops containing an appropriate dilution of fluorescently-conjugated secondary antibody (see below), overlaid with mineral oil and incubated in the dark at 4°C for 3 hours. A further three 70μl PBS-T washing steps were repeated before embryos were DNA counter-stained using Vectashield mounting media containing DAPI (Vector). Primary antibodies (and dilutions): a) raised in rabbit: i. anti-phospho-EIF4EBP1 (Thr37/46 – Cell Signalling Technologies # 9459; 1:50), ii. anti-pan-EIF4EBP1 (Cell Signalling Technologies # 9644; 1:50), iii. anti-phospho-EIF4EBP1 (Ser64 - Cell Signalling Technologies # 9451; 1:50), iv. anti-phospho-EIF4EBP1 (Thr70 - Cell Signalling Technologies # 9455: 1:50), v. anti-phospho-EIF4EBP1 (Thr37/46 - Cell Signalling Technologies # 2855: 1:200), 1:200), vi. anti-AMOT (kind gift of H. Sasaki; 1:100), vii. anti-PRKCZ (Santa Cruz Biotechnology #sc-216; 1:200), and viii. anti-PARD6B (Santa Cruz Biotechnology #sc-67393; 1:200), b) raised in mouse:, i. anti-CDX2 (Biogenex #MU392A-UC; 1:200) and ii. anti-YAP1 (Santa Cruz Biotechnology #sc-101199; 1:100). Secondary antibodies (and dilutions): i. donkey anti-rabbit-Alexa-Fluor^647^ (Abcam # ab150075; 1:500), ii. donkey anti-mouse-Alexa-Fluor^647^ (Abcam # ab150107; 1:500) and donkey anti-rabbit-Alexa-Fluor^488^ (Abcam # ab150073; 1:500). For F-actin counter-staining, immuno-fluorescent stained embryos were subject to additional processing, before the DAPI counterstain mounting in Vectashield, as follows; following the terminal three 70μl PBS-T washes (described above), embryos were transferred into 5μl drops, overlaid with mineral oil, of Oregon Green^488^ Phalloidin (O7466, ThermoFisher Scientific) diluted 1:50 in PBST and incubated at room temperature for 30 minutes (in the dark). Embryos were then washed through three 70μl drops of PBST and DNA counterstained in DAPI containing Vectashield, as described.

### Fixed sample confocal microscopy, image analysis, cell counting and statistics

Imaging protocols were as previously described [43]. Immuno-fluorescently and/or fluorescently labelled phalloidin stained embryos, of the desired developmental stage and experimental condition, were placed in small drops of PBS on the surface of glass microscope coverslip 35mm dishes (MatTek Corp.). Scanning confocal fluorescence imaging was conducted using an Olympus FLUOVIEW FV10i inverted confocal microscope, using experiment appropriate excitation wavelengths and emission detector settings. All embryos in comparable control and experimental groups were scanned with the same non-saturating imaging settings and exported in TIFF format for image analyses. Numbers of blastomeres were counted in FV10-ASW 4.2 Viewer software (Olympus) - note, for 16-cell stage analyses we only counted cells in embryos with exactly 16 cells and at the E3.5/32-cell stage we counted those ranging from 28-32 cells, to aid direct comparison. Fluorescence intensity was quantified in FIJI (freeware [85]), either as Corrected Total Cell Fluorescence (CTCF = Integrated Density – (Area of selected cell X Mean fluorescence of background readings)) or normalised for area of the quantified region in single z-stack or whole blastomere or embryo as specified in text. Intensity of fluorescence of membrane domains was quantified as average intensity across domain in a single z-stack with the largest length of the measured domain. All quantifications were normalised for background fluorescence. Statistical analysis was performed in R 4.2.1. (https://www.R-project.org/). Data normality was first tested by Shapiro-Wilk test and values were then compared by t-test (normal data distribution) or Mann-Whitney test (not normal data distribution).

### Time-lapse confocal microscopy live embryo imaging and analyses

Post-microinjected 2-cell stage embryos (expressing recombinant fluorescent mRNAs – see above) were cultured to compacted 8-cell stage (E2.5+4h) and transferred into KSOM+AA imaging plates (Caisson Laboratories) supplemented with Torin1 (Selleckchem, 20µM) dissolved in DMSO (Merck, Czech Republic) or DMSO only. Complete embryo z-series time-lapse imaging was performed on a Leica SP5 confocal microscope, equipped with EMBL incubator set to 5% CO_2_ at 37°C. The wavelengths 458nm, 514nm and 561nm were used for excitation, HCX PL APO CS 40× water objective NA 1.1 and HyD detectors were used for detection of CFP, Venus and mCherry signal, respectively. 41 z-stacks were taken every 15 minutes for each single embryo position. Pre-M-phase 8-cell nuclei position along radial axes and mitotic spindle angles were measured in IMARIS 6.2.1 (BitPlane). For nuclear positioning, the distance between apical domain and nucleus surface, nucleus diameter, and the distance between nucleus surface and basal domain were measured from time frame images immediately preceding nuclear envelope breakdown. Mitotic spindle angles were quantified using the coordinates of the centre of the embryo and spindle poles, immediately prior to anaphase onset.

### O-propargyl-puromycin (OPP) quantification of *de novo* protein synthesis

OPP staining was performed using a Click-iT Plus OPP Alexa Fluor 488 Protein Synthesis Assay Kit (ThermoFisher Scientific; Cat. No. C10456). Mouse embryos were first cultured *in vitro* from E1.5 to E2.5+7h in KSOM+AA as described above, and then transferred to KSOM+AA supplemented with 20 µM Torin1 dissolved in DMSO, or equivalent volume of DMSO only, and 5µM OPP Reagent, where they were incubated for 10 minutes. The embryos were then immediately fixed, permeabilized, and washed as described above. Click-iT reactions were set up according to the kit manual. The fixed embryos were incubated in the reaction mixture for 25 minutes at room temperature in the dark. Thereafter, the embryos were washed in a 1:1 mixture of kit-provided wash buffer and PBST, DNA counterstained in DAPI containing Vectashield, and imaged on confocal microscope as described above.

### Mass spectrometry

Employed protocols to survey the general proteome by mass spectrometry in preimplantation mouse embryos +DMSO (control) or +mTORi (+Torin1), 3 biological replicates each, were conducted as previously described [60]. Briefly, embryos at the desired developmental stage during 8-to 16-cell transition (E2.5+8 and E2.5+9 hours) were lysed in 5µl of SDT buffer (4% SDS, 0.1M DTT, 0.1M Tris/HCl pH 7.6), incubated at 95°C for 12 min and frozen at -80°C until further processing. Individual protein solutions were processed using filter-aided sample preparation (FASP) method as described previously [60]. FASP eluates were transferred into the LC-MS vials and analyzed using Ultimate 3000 RSLCnano system connected to Orbitrap Fusion Lumos Tribrid mass spectrometer (Thermo Fisher Scientific) as described previously [60] with the following changes. Peptides trapped on the trap column were eluted and separated on the analytical column using 130 minute long nonlinear gradient program (1-56% of mobile phase B; mobile phase A: 0.1% FA in water; mobile phase B: 0.1% FA in 80% ACN; start at 1% B, 95min 30% B, 130min 56 % B). Mass spectrometry data were acquired in a data-dependent strategy with top 20 approach and with survey scan (350-2000 m/z). The resolution of the survey scan was 120,000 (at *m/z* 200) with a target value of 4×10^5^ ions and maximum injection time of 100 ms. HCD MS/MS (30% relative fragmentation energy) spectra were acquired with a target value of 5.0×10^4^. The MS/MS spectra were recorded in Orbitrap at resolving power of 15000 (200 m/z) and the maximum injection time for MS/MS 22 ms. Dynamic exclusion was enabled for 30 seconds after one MS/MS spectrum acquisition. The isolation window for MS/MS fragmentation was set to 1.2 m/z. The analysis of the mass spectrometric RAW data files was carried out using the MaxQuant software (version 1.6.2.10) using default settings unless otherwise noted. MS/MS data searches were done against modified cRAP database (based on http://www.thegpm.org/crap, 112 protein sequences) containing protein contaminants like keratin, trypsin etc., and UniProtKB protein database for *Mus musculus* (https://ftp.uniprot.org/pub/databases/uniprot/current_release/knowledgebase/reference_proteomes/Eukaryota/UP000000589/UP000000589_10090.fasta.gz; downloaded 2019-05-08, version 2019/05, number of protein sequences: 22,287). Oxidation of methionine and proline, deamidation (N, Q) and acetylation (protein N-terminus) as optional modification, and trypsin/P enzyme with two allowed miss cleavages and minimal peptide length of 6 amino acids were set. Peptides and proteins with FDR threshold <0.01 and proteins having at least one unique or razor peptide were considered only. Match between runs was set for all analyses and second peptides option was checked. Protein intensities reported in proteinGroups.txt file and evidence intensities reported in evidence.txt file (output of MaxQuant program) were further processed using the software container environment (https://github.com/OmicsWorkflows), version 3.7.1a. Processing workflow is available upon request. Briefly, it covered: a) removal of decoy hits and contaminant protein groups, b) protein group intensities log_2_ transformation, c) LoessF normalization and d) differential expression using LIMMA statistical test (qualitative changes were considered separately without statistical evaluation). Protein candidates were selected based on the following criteria: statistically significant difference (p-value<0.05) in at least one of the timepoints and biological relevance based on published literature.

### Quantitative RT-PCR

Quantitative RTPCR (Q-RTPCR) was performed essentially as described [43]. Total RNA was prepared from ∼30 cultured 16-cell stage (E3.15) embryos that had been microinjected with gene specific or GFP/NTC control dsRNA/siRNA (as described above – i.e. in both blastomeres of late 2-cell stage embryos) and total RNA purified as instructed (Arcturus Biosciences; ‘PicoPure RNA isolation’). Eluted RNA (10μl) was DNaseI treated (Ambion; ‘DNA-free’ kit) and used to derive cDNA (30μl) using oligo-dT priming (Invitrogen; ‘SuperscriptIII Reverse Transcriptase’). 0.5μl of diluted cDNA (1:3 - nuclease-free water) was used as template in 10μl real-time PCR reactions (Qiagen: ‘SYBR Green PCR kit’) to assay specific transcripts (BioRad, ‘CFX96 Real-Time System’). Gene transcript specific oligonucleotide primer sequences used (final reaction conc. 400nM): *Eif4g* (s - ACCCATGGGCAAAGCTACT, a - ACAGCATCCCCACCTTTTT), *Larp1* (s - CTCGACCCTCACCAGCAC, a - GCTCATCCTGATCCTTAGACATC), *Ank2* (s - TGAGAGTCTGCCACCTGTTG, a - TGCTCATCTTGGGGATCTTC), *Dctn2* (s - TCTGGGACCAGATGCTGCAA, a - TCAGGCCGTGAGTGGAGTTC), *Ddx21* (s - TTCCTTCTGCAACGGAAATAA, a - GAGGCACAGAATCCAAGAGC) and *H2afz* (s - GCGCAGCCATCCTGGAGTA, a - CCGATCAGCGATTTGTGGA). Transcript levels were internally normalised against *H2afz* (encoding histone H2A) levels, and fold changes (plus s.e.m.) after dsRNA/siRNA mediated knockdown derived using the ΔΔCt method [86]. A minimum of two biological replicates of at least three technical replicates were employed.

## ACKNOWLEDGEMENTS

We acknowledge the Institute of Parasitology (Biology Centre of the Czech Academy of Sciences, in České Budějovice) for housing mice, Marta Gajewska (Institute of Oncology, Warsaw, Poland) and Anna Piliszek (Institute of Genetics and Animal Breeding, Polish Academy of Sciences, Jastrzębiec, Poland) for founder CBA/W mice and other members of our laboratory for valuable inputs and discussions. CIISB (Instruct-CZ Centre of Instruct-ERIC EU consortium) funded by MEYS CR infrastructure project LM2023042 is gratefully acknowledged for the financial support of the measurements at the CEITEC Proteomics Core Facility. Computational resources for mass spectrometry data were provided by the e-INFRA CZ project (ID:90140), supported by the Ministry of Education, Youth and Sports of the Czech Republic. The anti-AMOT antibody used in IF was a generous gift from Hiroshi Sasaki (Osaka University, Japan).

## AUTHOR CONTRIBUTIONS

The project was devised and supervised by AWB, LG & AS. All characterisation of mTORi/p38-MAPKi embryo phenotypes (including specific RNAi mediated gene expression knockdowns and IF) were performed by LG (as lead investigator), JT, PC, PB, MK and GV. Live confocal microscopy cell imaging was conducted by KK and MA. Mass spectrometry completed by DP, PH & ZZ. AWB and LG wrote the manuscript (in cooperation with all other authors).

## DATA AVAILABILITY

The mass spectrometry proteomics data are deposited to the ProteomeXchange Consortium via the PRIDE partner repository with the dataset identifier PXD039423. Reviewer login information are as follows, user, password: Xfvb1gXr.

## FIGURE LEGENDS

## SUPPLEMENTARY FIGURE & TABLE LEGENDS

**Figure S1:**
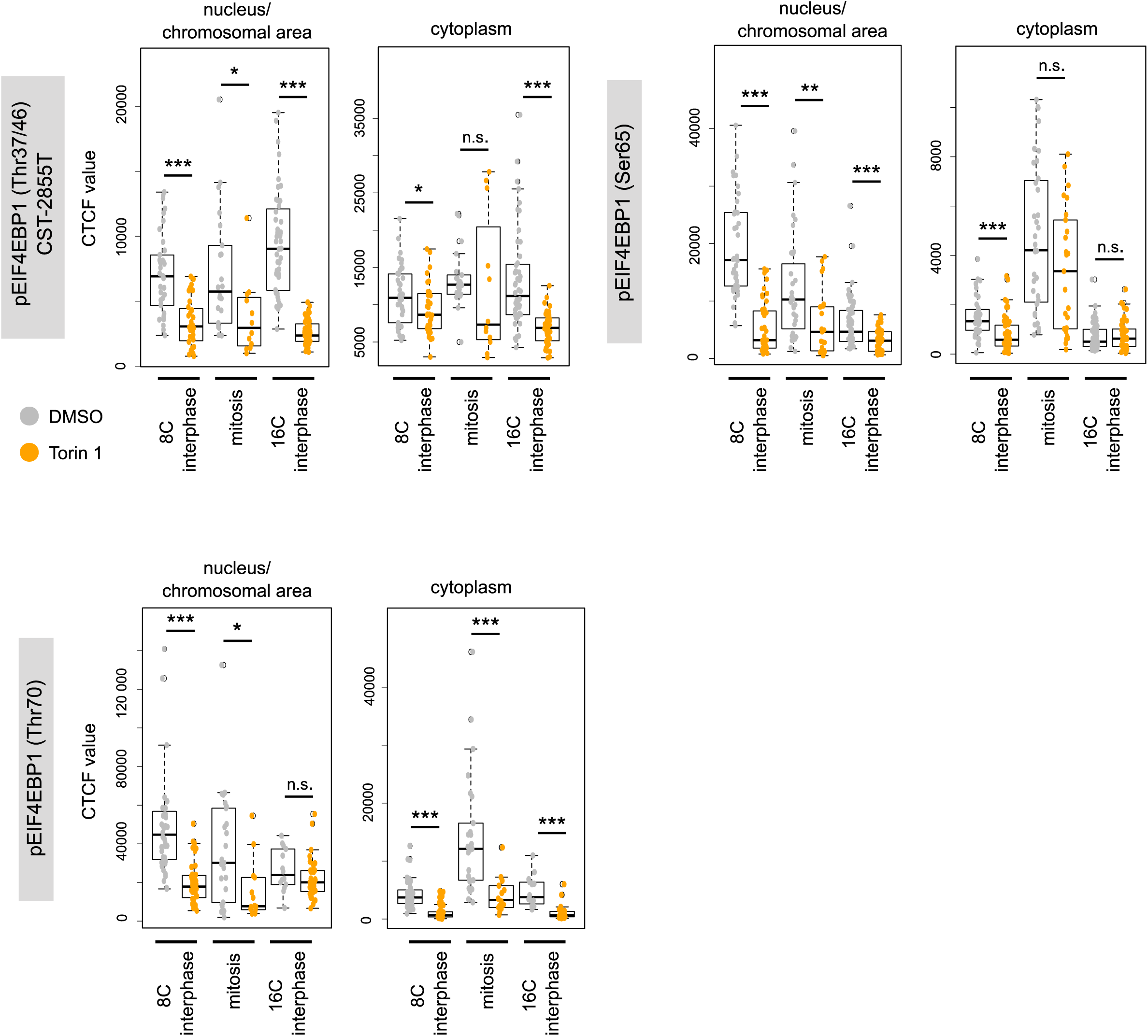
mTOR-regulated cap-dependent translation occurs during 8- to 16-cell division (supplementary). Quantification of individual cell IF staining assays using additional antibodies recognising pEIF4EBP1 (Thr37/46), pEIF4EBP1 (Ser65) and pEIF4EBP1 (Thr70) during either 8- or 16-cell stage interphases and in dividing 8-cell stage blastomeres transiting to the 16-cell stage, +/-Torin1.

**Figure S2:**
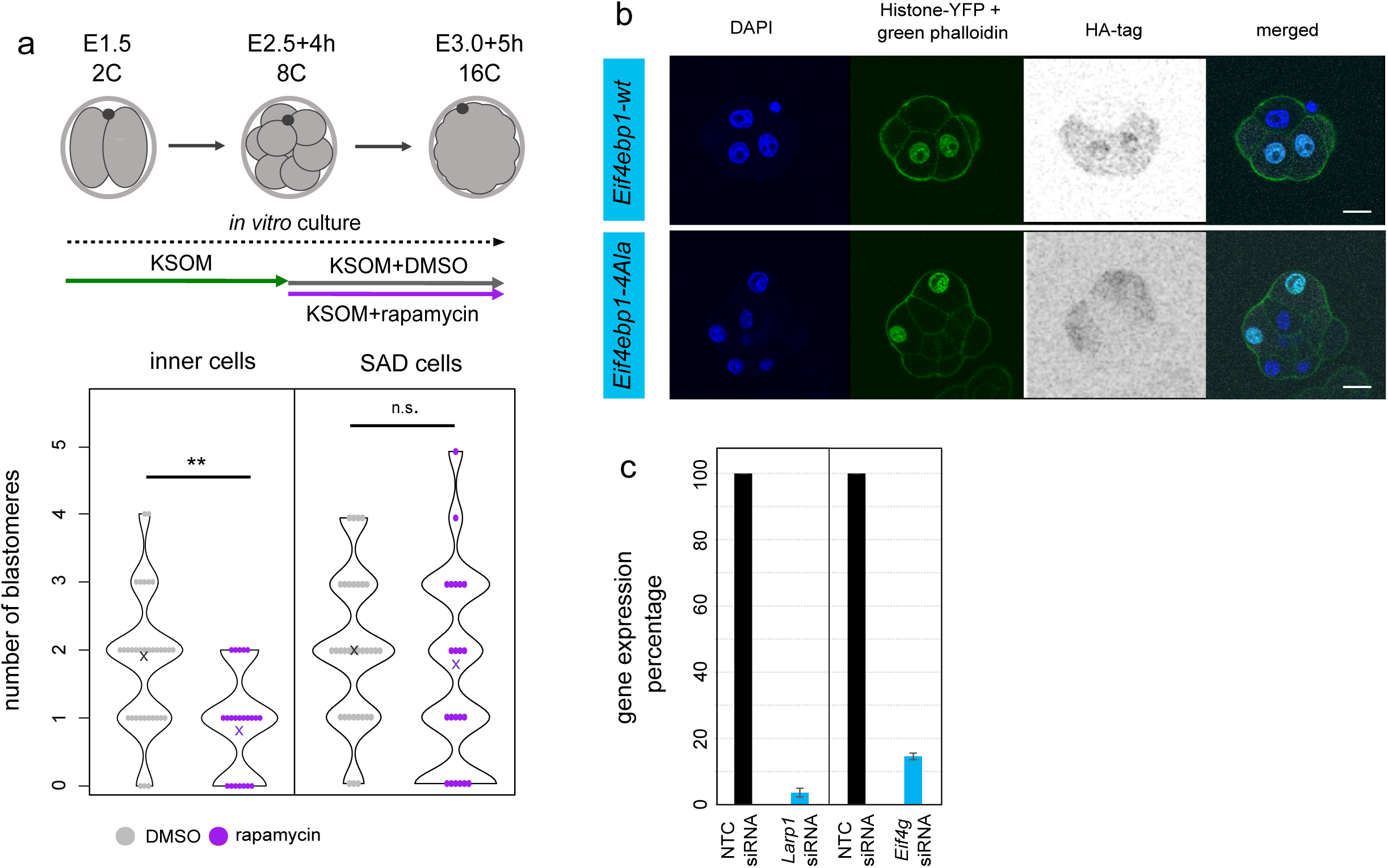
mTOR-regulated 7mG-cap-dependent translation plays a role in the formation of primary ICM founder cells (supplementary). **a)** Quantification of the number of inner and SAD cells in 16-cell embryos +/-Rapamycin, “x” marks an average value. **b)** IF staining confirming the expression of microinjected mRNA constructs encoding HA-tagged EIF4EBP1-wt and EIF4EBP1-4Ala (greyscale). Scale: 20µm. **c)** Confirmation of efficient *Larp1* and *Eif4g* mRNA downregulation by quantitative RT-PCR in whole 16-cell embryos after injecting both blastomeres at 2-cell stage with gene-specific siRNAs vs NTC/control siRNA.

**Figure S3:**
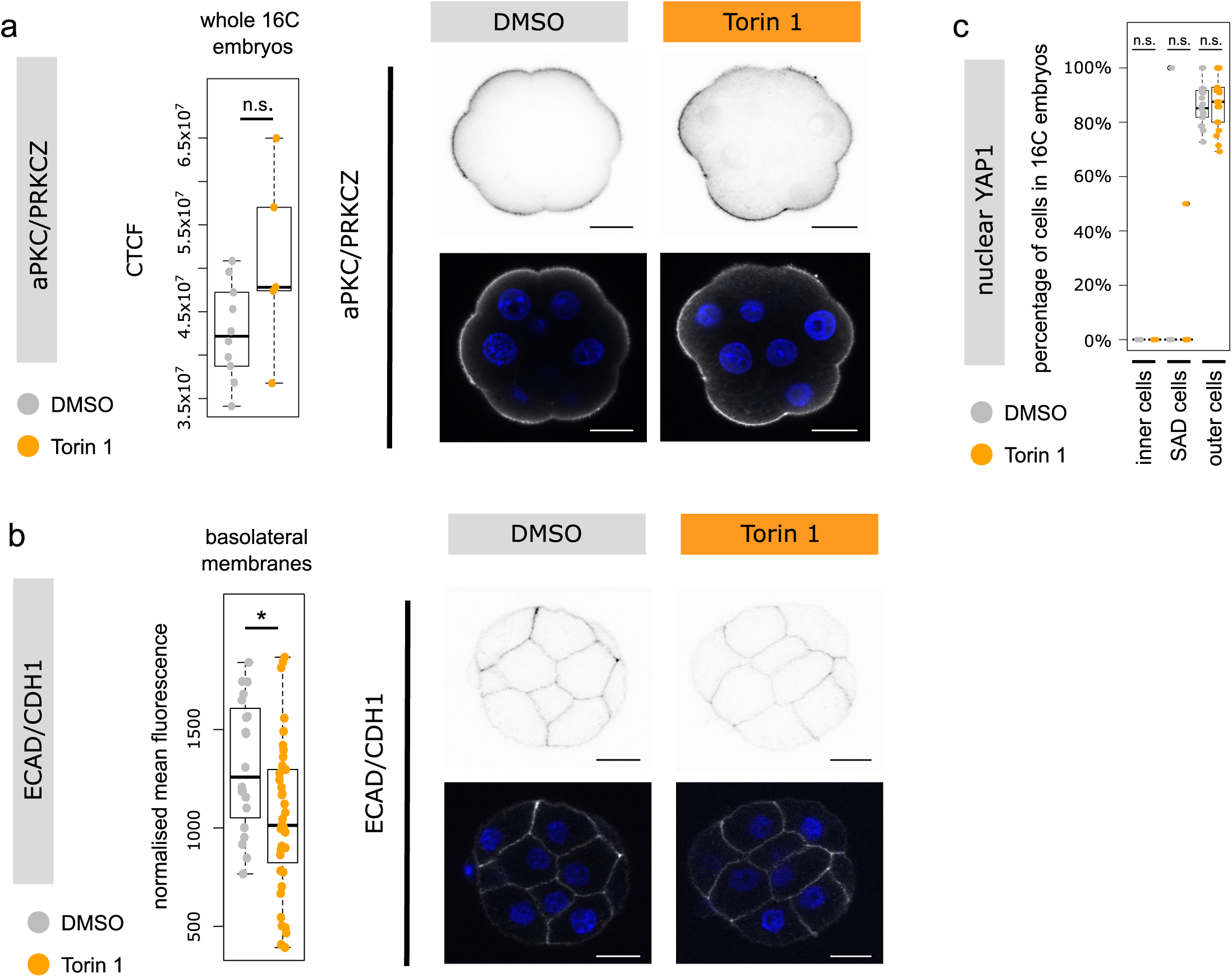
16-cell stage apical-basolateral polarisation is not affected under mTORi conditions. Quantification of normalised IF fluorescence intensity and representative IF micrographs for assays recognising **a)** aPKC/PKRCZ (single confocal z-slice shown), **b)** ECAD/CDH1 (single confocal z-slice shown) and **c)** Quantification of the proportion of blastomeres with nuclear YAP1 IF staining, +/-Torin1.

**Figure S4:**
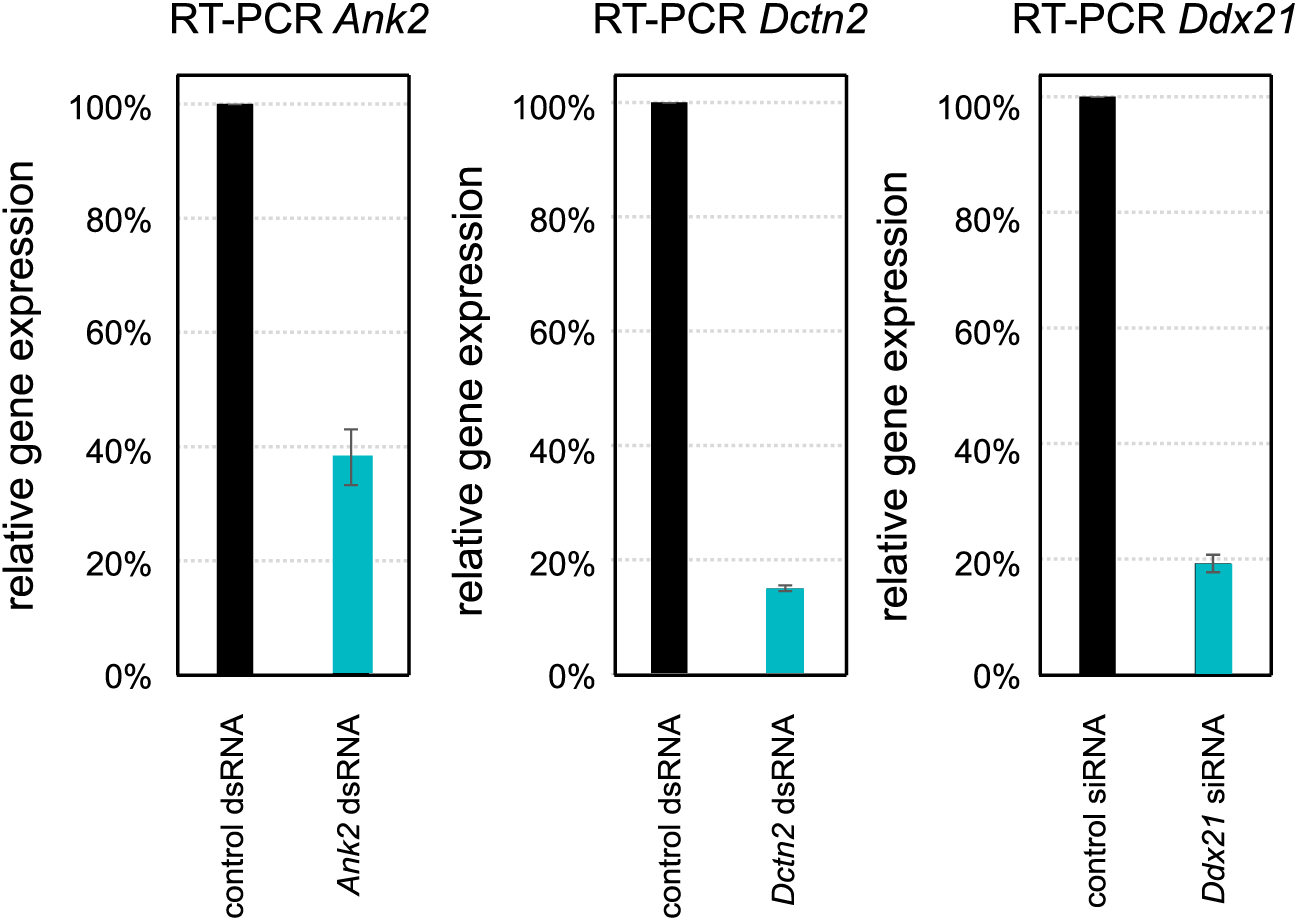
Confirmed RNAi mediated expression knockdown of candidate TOP-motif containing mRNAs. Relative mRNA expression level of *Ank2, Dctn2* and *Ddx21* transcripts as assayed by quantitative RT-PCR in whole 16-cell embryos after injecting both blastomeres at 2-cell stage with gene-specific dsRNAs/siRNA or control dsRNA/siRNAs (expression set as 100%); dsRNAs targeting *Ank2*, *Dcnt2* and *GFP* (control) transcripts and siRNAs against *Ddx21* (plus NTC control).

**Figure S5:**
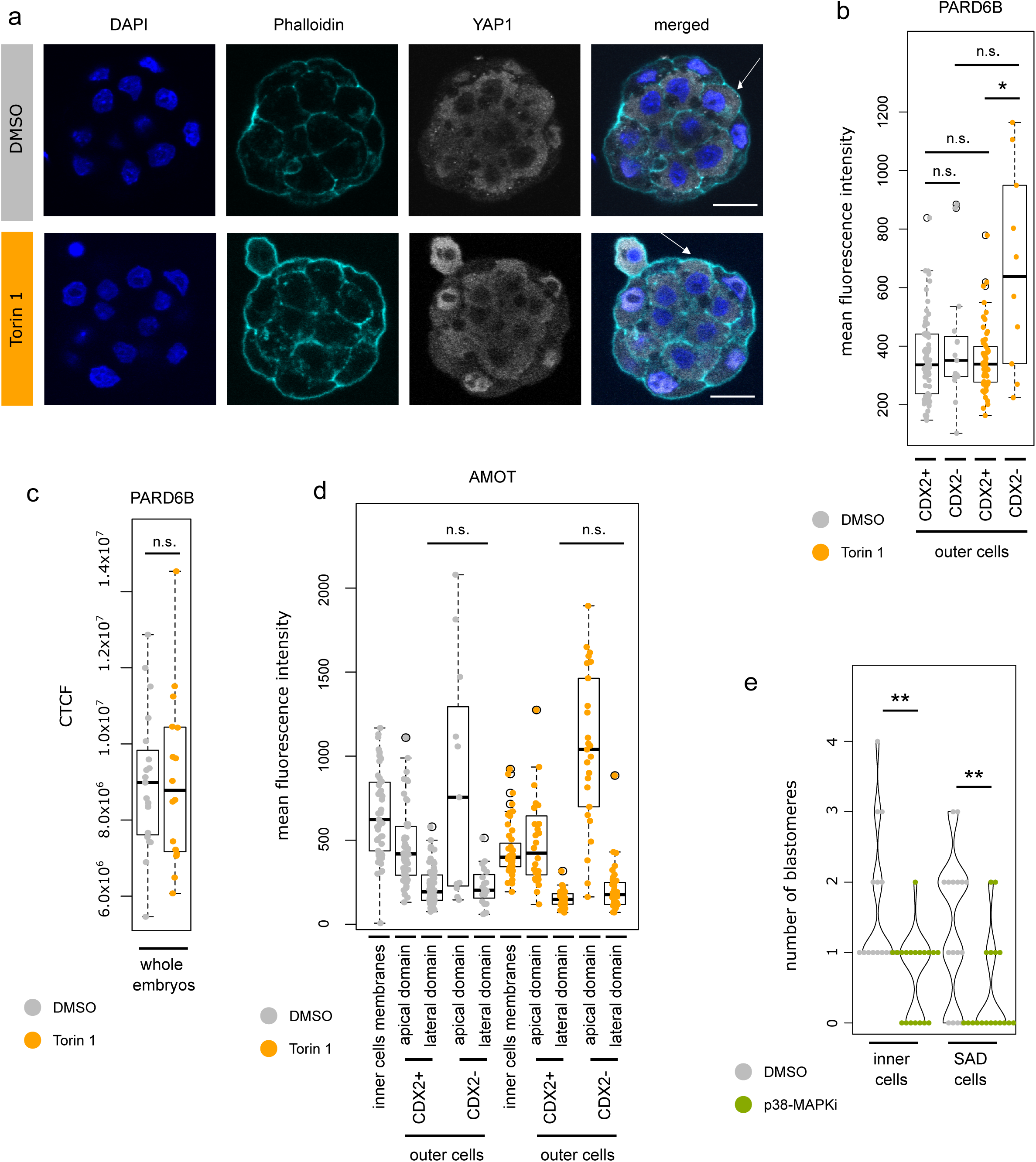
Supernumerary outer cells in mTORi 32-cell stage embryos exhibit molecular characteristics of ICM-like cells (supplementary). **a)** Example IF staining micrographs of YAP1 staining in 32-cell stage embryos cultured under control and mTORi conditions. YAP1 visualised in greyscale, DAPI in blue, phalloidin in cyan, representative confocal z-stacks shown. Scale: 20µm. Example outer blastomeres with cytoplasmic YAP1 and various levels of nuclear YAP1 are marked by an arrow. **b)** Quantification of mean apical membrane IF fluorescence intensity of PARD6B in single z-stack (with largest membrane length) in individual outer cells +/-CDX2, +/-Torin1. **c)** Quantification of whole embryo IF fluorescence intensity of PARD6B +/-Torin1. **d)** Quantification of basal membrane AMOT mean IF fluorescence intensity in single z-stack (with largest membrane length) in individual inner cells and outer cells +/-CDX2, +/-Torin1. **e)** Quantification of the number of inner and SAD cells in 16-cell embryos, +/-p38-MAPKi.

**Table S1:**
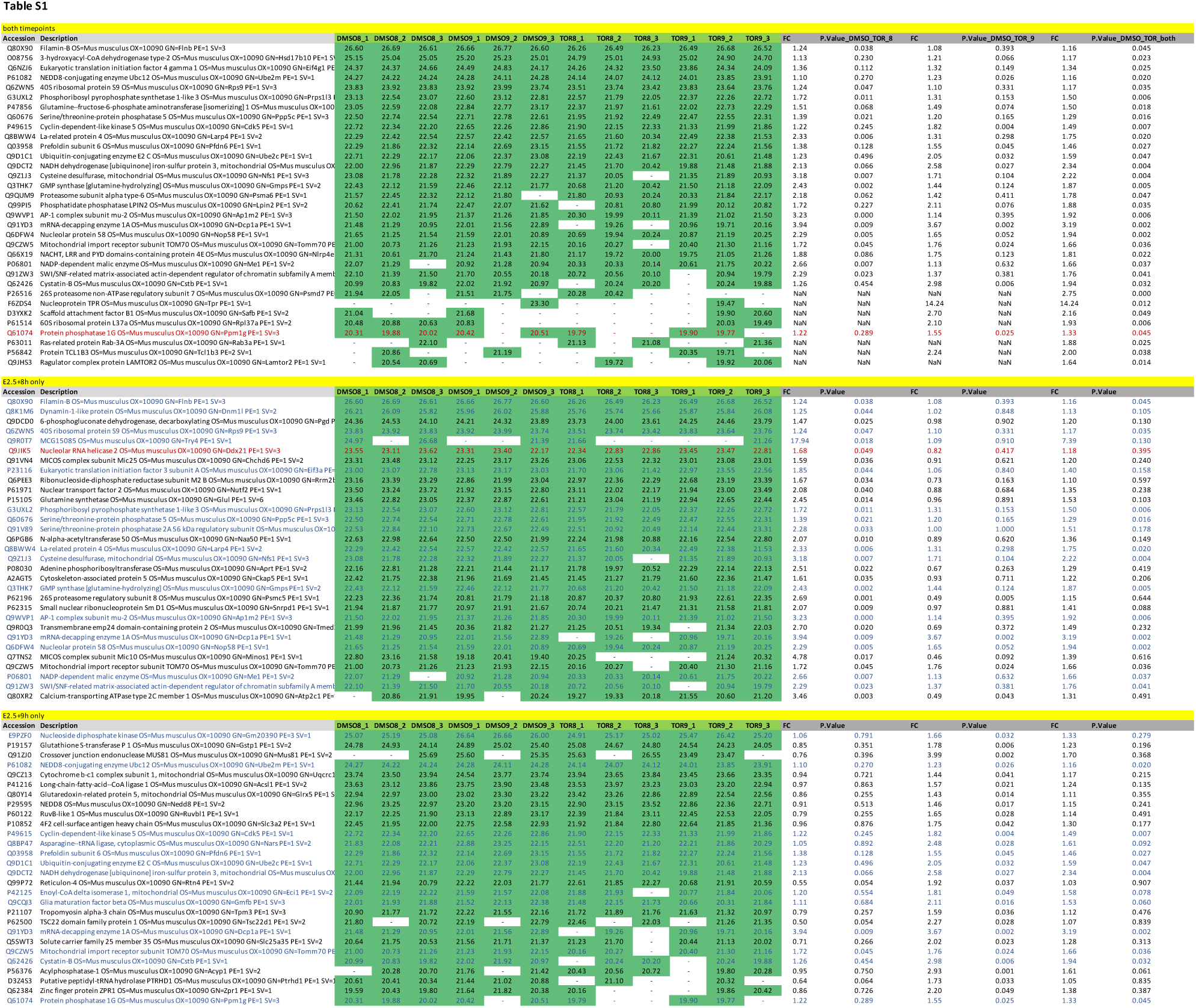
List of significantly differently changing proteins between control and mTORi embryos at E2.5+8h and E2.5+9h.

